# A mini-TGA protein, lacking a functional DNA-binding domain, modulates gene expression through heterogeneous association with transcription factors

**DOI:** 10.1101/2022.03.30.486221

**Authors:** Špela Tomaž, Marko Petek, Tjaša Lukan, Karmen Pogačar, Katja Stare, Erica Teixeira Prates, Daniel A. Jacobson, Jan Zrimec, Gregor Bajc, Matej Butala, Maruša Pompe Novak, Quentin Dudley, Nicola Patron, Ajda Taler-Verčič, Aleksandra Usenik, Dušan Turk, Salomé Prat, Anna Coll, Kristina Gruden

## Abstract

TGA transcription factors, which bind their target DNA through a conserved basic region leucine zipper (bZIP) domain, are vital regulators of gene expression in salicylic acid (SA)-mediated plant immunity. Here, we investigate the role of StTGA2.1, a potato TGA lacking the full bZIP, which we name a mini-TGA. Such truncated proteins have been widely assigned as loss-of-function mutants. We, however, confirm that *StTGA2.1* overexpression compensates for SA-deficiency. To understand the underlying mechanisms, we show that StTGA2.1 can physically interact with StTGA2.2 and StTGA2.3, while its interaction with DNA was not detected. We investigate the changes in transcriptional regulation due to *StTGA2.1* overexpression, identifying direct and indirect target genes. Using *in planta* transactivation assays, we confirm that StTGA2.1 interacts with StTGA2.3 to activate *StPRX07*, a member of class III peroxidases, which are known to play role in immune response. Finally, via structural modelling and molecular dynamics simulations, we hypothesise that the compact molecular architecture of StTGA2.1 distorts DNA conformation upon heterodimer binding to enable transcriptional activation. This study demonstrates how protein truncation can lead to novel functions and that such events should be studied carefully in other protein families.

## INTRODUCTION

Plants have developed efficient strategies to withstand the invasion of surrounding microbes. Pathogen recognition is mediated by plant cell-surface and intracellular receptors, triggering a cascade of intracellular reactions, orchestrated by phytohormones, ultimately leading to a finely modulated transcriptional reprogramming^1^. Regulation of defence-related gene expression is among the most fundamental aspects of the immune response, involving multiple transcription factors and cofactor proteins. Since their initial discovery in tobacco over 30 years ago^2^, the importance of TGACG-binding (TGA) transcription factors in plant immunity, as well as modulation of other cellular processes, has been widely studied^3^.

TGAs are a group of transcription factors belonging to the basic region leucine zipper (bZIP) protein family. Their mechanism of action has been thoroughly studied in *Arabidopsis thaliana*, where the ten Arabidopsis TGAs (AtTGAs) group into five clades^4^. Clade II members, AtTGA2, AtTGA5, and AtTGA6, are essential regulators of the salicylic acid (SA)-mediated defence response, where they play a redundant, yet vital role in establishing resistance following infection^1,5^. They co-regulate the expression of key defence-related genes and genes involved in SA synthesis through interaction with NON-EXPRESSOR OF PR (NPR) cofactors^6,7^, while also participating in jasmonic acid and ethylene-mediated signalling^8^. Structurally, TGAs consist of an intrinsically disordered N-terminus of varying length, a conserved bZIP domain, which entails a basic region and a leucine zipper, and a C-terminal region that contains a putative Delay of Germination 1 (DOG1) domain (*under review, Š. T., K.G. and A.C.*). TGAs bind their target DNA through the bZIP basic region, while the leucine zipper is important for protein dimerization^9^ and oligomerization^10^. The TGACG core sequence is sufficient for TGA binding, although high-throughput DNA-binding studies revealed the TGACGTCA palindrome as the representative binding motif^11,12^.

The molecular mechanisms of TGA-mediated regulation involve complex interactions between TGAs and other proteins^3^. For example, AtTGA2 acts as a constitutive repressor of defence-related *PR-1* gene expression in absence of biotic stress^5,13,14^, whereas AtTGA6 has been shown to activate *PR-1* in absence of AtTGA2^13^. The repressive activity of AtTGA2 is alleviated by the SA-receptor AtNPR1, which affects binding stoichiometry through interaction with AtTGA2 N-terminus, leading to the formation of a transcriptional activation complex^10,14^. Other regulators, such as WRKY50^15^ and histone acyltransferase (HAC) transcription factors^16^, have also been shown to contribute to AtTGA2 transcriptional function.

Although the results obtained in Arabidopsis provide a molecular framework for understanding the role of TGAs in plant immunity, we know much less about their function in crops. The involvement of TGAs in biotic stress response has been reported in several species, including rice^17^, soybean^18^, strawberry^19^, tobacco^9^, and tomato^20^. Potato (*Solanum tuberosum* L.) is one of the most widely grown crops^21^ and tuber production is severely threatened by pathogen infections. Several transcription factor families have been associated with the regulation of potato defence response^22^, but the mechanisms underlying potato TGA (StTGA) activity remain largely unexplored.

Here we identify the mini-TGA StTGA2.1, a potato clade II TGA, which lacks most of the bZIP DNA-binding domain and has a shorter N-terminus. We hypothesize that StTGA2.1 cannot bind DNA by itself because of the truncated bZIP and therefore modulates gene expression through its interaction with additional DNA-binding StTGAs. By combining *in vivo* and *in vitro* functional studies, we confirm the role of StTGA2.1 in potato immunity. Furthermore, using *in silico* structural analysis and molecular dynamics simulations, we provide insights into the molecular basis for a different mechanism of action of StTGA2.1 compared to other StTGAs.

## RESULTS

### Potato encodes clade II TGAs with truncated bZIP domain

In previous work, we investigated gene expression in response to viral infection in non-transgenic resistant potato (NT) and its transgenic derivative (NahG), which is impaired in SA accumulation and thus sensitive to infection^23^. To identify the TGA transcription factors involved in potato immunity, we examined the expression patterns of the fourteen StTGA genes, orthologues of AtTGAs (Supplementary Tables 1,2). Notably, *Sotub10g022560* was up-regulated in infected NahG transgenic plants, but not in the parental lines, suggesting that it may be an important component of SA signalling.

To classify the StTGAs, we conducted a phylogenetic analysis of all candidate potato proteins, along with the ten AtTGAs and thirteen TGAs from tomato (SlTGAs)^24,25^. Interestingly, the three clade II AtTGAs are orthologous to five StTGAs and four SlTGAs (Fig. 1a). Three closely related members of this clade, including *Sotub10g022560*, named StTGA2.1, StTGA2.4 (*Sotub10g022570*) and SlTGA2.3 (*Solyc10g080780*)^24,25^ have shorter protein sequences than other TGAs (Fig. 1b). Domain prediction studies showed that they retain the putative C-terminal DOG1 domain, however, the bZIP domain is almost completely lost, retaining only a partial zipper region. In addition, their N-terminus is very short and dissimilar to the N-termini of other clade II TGAs. We named these three proteins mini-TGAs.

**Fig. 1.**
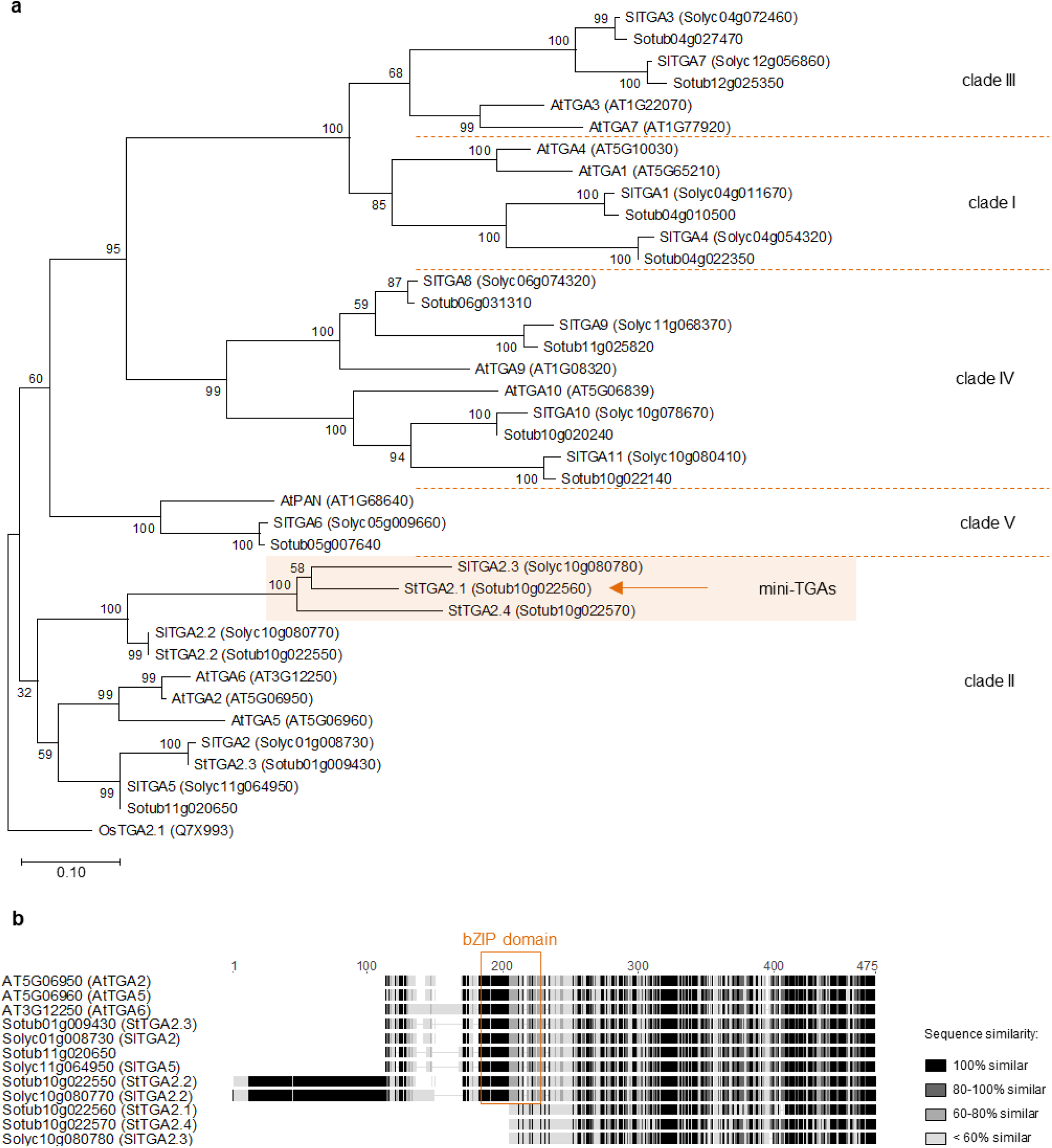
Phylogenetic analysis and domain characterization of StTGAs. **a,** A rooted phylogenetic tree of potato, tomato and Arabidopsis TGAs. The mini-TGA branch is shaded in orange and StTGA2.1 is marked (arrow). The branch length scale represents the number of amino acid substitutions per site. The rice OsTGA2.1^26^ serves as tree root. **b,** Protein sequence alignment of clade II TGAs, showing the position of the bZIP domain (orange box) and the shorter sequences of mini-TGA members, StTGA2.1, StTGA2.4 and SlTGA2.3. The alignment is coloured with the Geneious Prime (https://www.geneious.com) sequence similarity colour scheme, based on the identity score matrix. Sequence numbering (aa) is shown above the alignment.

By targeted sequencing of a ∼36.5 kbp region on chromosome 10, where *StTGA2.1*, *StTGA2.2* (*Sotub10g022550*) and *StTGA2.4* loci are co-located, we confirmed the reduced length of *StTGA2.1* and *StTGA2.4* in a tetraploid cultivar that was used for further analyses (Supplementary Fig. 1).

### StTGA2.1 improves immune response in salicylic acid-deficient potato

SA signalling has proven vital for the establishment of an efficient defence response against potato virus Y (PVY) infection in resistant potato cultivars^23,27^. We thus investigated the role of StTGA2.1 in plant immunity using the potato-PVY pathosystem. We generated SA-deficient NahG transgenic potato plants inducibly overexpressing *StTGA2.1* (TGA2.1-NahG) using the glucocorticoid-system^28^, in which target gene expression is controlled by external application of dexamethasone (DEX). Two transgenic TGA2.1-NahG lines, showing more than 6-fold induction in *StTGA2.1* expression after DEX treatment (Supplementary Fig. 2a,b), were selected for further analysis. We observed that viral replication was significantly reduced in TGA2.1-NahG compared to NahG at 10 days post infection (dpi) (Fig. 2, Supplementary Fig. 2c-e). As expected, little to no PVY was detected in NT plants exhibiting a typical resistant phenotype^23^. This shows that overexpression of *StTGA2.1* can compensate for the lack of SA in potato immune response to PVY.

**Fig. 2.**
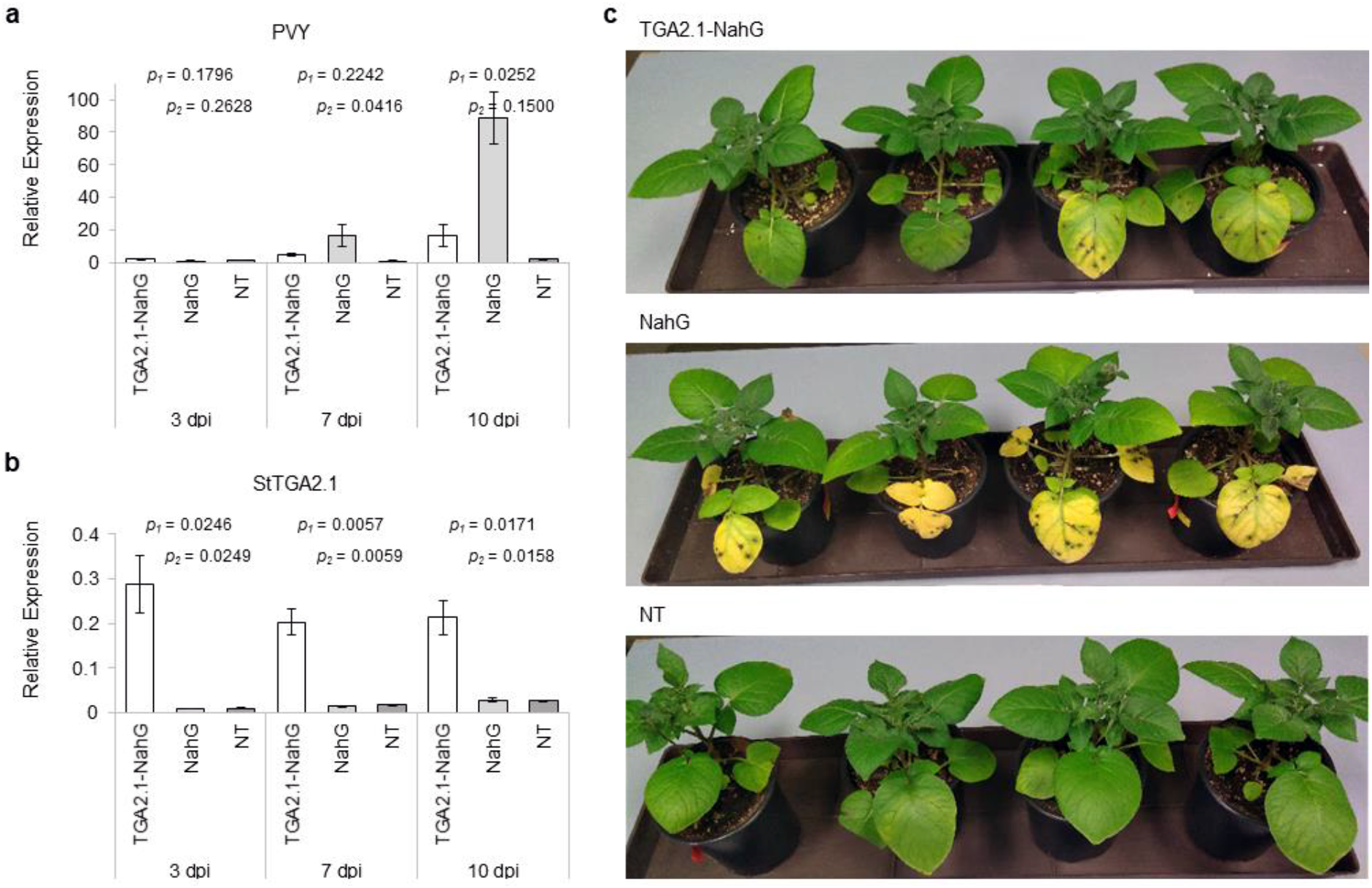
StTGA2.1 attenuates PVY replication in salicylic acid-deficient plants. Relative expression levels of **a,** PVY and **b,** *StTGA2.1* in PVY-infected leaves of dexamethasone (DEX)-treated TGA2.1-NahG (white), NahG (light grey) and NT (dark grey) plants at 3, 7 and 10 days post infection (dpi). Average values ± standard error from three biological replicates are shown. Significance was determined using a two-tailed *t*-test comparing TGA2.1-NahG with NahG (*p_1_*) and TGA2.1-NahG with NT (*p_2_*). **c,** Phenotypic differences in PVY-infected leaves of DEX-treated TGA2.1-NahG, NahG and NT plants at 10 dpi.

### StTGA2.1 retains its dimerization ability and shows a distinct localization pattern

Protein interaction studies using the yeast two-hybrid assay showed that StTGA2.1 can form both homodimers and heterodimers with StTGA2.2 and StTGA2.3 (*Sotub01g009430*) (Fig. 3a), further confirmed by *in planta* co-immunoprecipitation assay (Fig. 3b, Supplementary Fig. 3a,b). Additionally, the size-exclusion chromatography elution volume of a recombinant His_6_-tagged StTGA2.1 corresponded to the size of a dimer (Supplementary Fig. 3c,d), while chemical cross-linking of a non-tagged protein yielded monomers, dimers, and higher order complexes (Supplementary Fig. 3e). Overall, these results demonstrate that StTGA2.1 retains protein-protein interaction ability. In addition, we examined whether StTGA2.1 can interact with two potato NPR cofactors, an orthologue of AtNPR1, StNPR1 (*Sotub07g011600*), and an orthologue of AtNPR3 and AtNPR4, StNPR3/4 (*Sotub02g015550*). Our results showed that StTGA2.1 as well as StTGA2.2 and StTGA2.3 interact with both StNPRs in yeast and that the addition of SA to the media promotes these interactions (Supplementary Fig. 4). Thus, the ability to interact with NPR proteins is not perturbed in mini-TGA StTGA2.1.

**Fig. 3.**
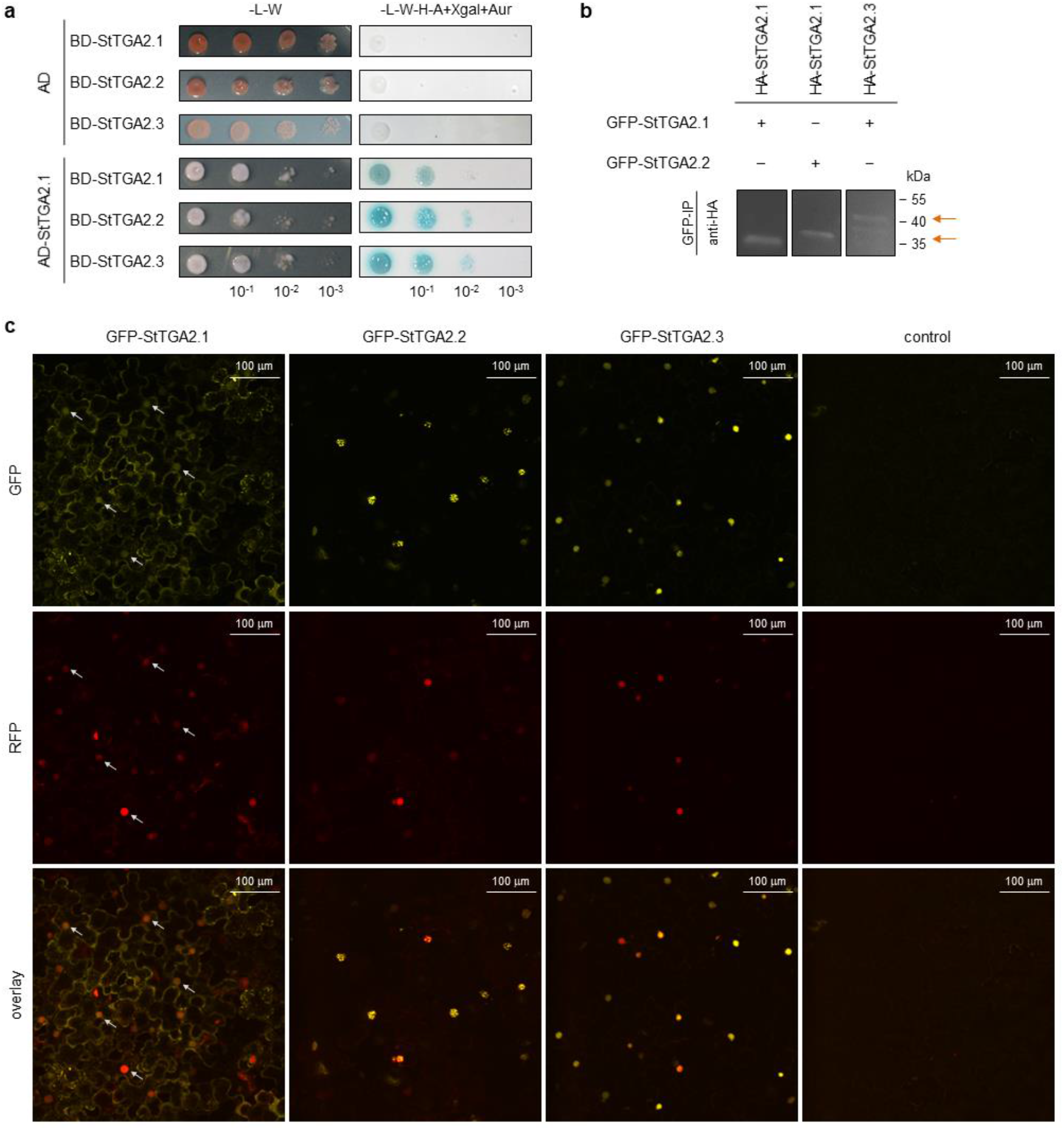
StTGA2.1 can form homodimers, heterodimers and localizes to diverse cellular compartments. **a**, StTGA2.1 interactions with itself, StTGA2.2, or StTGA2.3 in the yeast two-hybrid assay. Yeast were co-transformed with bait (BD) and prey (AD) construct combinations and selected on control media without Leu and Trp (-L-W). Positive interactions were determined by yeast growth on selection media without Leu, Trp, His and adenine, with added X-α-galactosidase and Aureobasidin A (-L-W-H-A+Xgal+Aur). **b,** StTGA2.1 interactions with itself, StTGA2.2, or StTGA2.3 in the co-immunoprecipitation assay. The combination of GFP and HA-tagged proteins expressed in *N. benthamiana* is indicated for each sample (+/−). Positive interactions were determined by detection of immunoprecipitated (GFP-IP) complexes with anti-HA antibodies. Arrows indicate expected bands. Controls are shown in Supplementary Fig. 3. **c,** Subcellular localization of GFP-tagged StTGA2.1, StTGA2.2 and StTGA2.3 (yellow) with H2B-RFP nuclear marker (red) in *N. benthamiana* leaves. The *p19* silencing suppressor was expressed as control. Protein fluorescence is represented as the z-stack maximum projection. Arrows indicate examples of StTGA2.1 nuclear localization. Scale bars, 100 μm.

Subcellular localization of GFP-tagged StTGA2.1 in the *N. benthamiana* leaf epidermis and mesophyll showed that it can localize to cell nuclei (Fig. 3c). StTGA2.1 also showed a distinct localization pattern with intense fluorescence in the cytoplasm, which was enhanced around the chloroplasts (Supplementary Fig. 5a). We also detected its fluorescence in the ER and in granular formations of about 0.5-1.0 μm in size (Supplementary Fig. 5). In contrast, StTGA2.2 and StTGA2.3 showed predominantly nuclear localization and were organized into subnuclear formations of different sizes within the nuclei (Fig. 3c, Supplementary Fig. 6).

### Identification of potential StTGA2.1 targets with spatial transcriptomic profiling

To gain insight into the mini-TGA mechanism of action in plant immunity, we examined the expression profile of plants overexpressing *StTGA2.1*. By sampling tissue sections containing lesions and their immediate surrounding area after PVY infection (Supplementary Fig. 7a), we were able to follow transcriptomic changes in PVY-responding cells^27^. RNA sequencing results showed a regulation of 217 genes due to *StTGA2.1* overexpression (TGA2.1-NahG vs. NahG plants comparison, GEO accession GSE196078). However, over 1,800 genes were differentially expressed exclusively in TGA2.1-NahG, when plants were exposed to pathogen infection (Supplementary Fig. 7b). Technical validation of the RNA sequencing data by qPCR is shown in Supplementary Table 3.

Gene Set Enrichment Analysis (Supplementary Table 4) enabled us to extract key differences in gene expression at the level of processes or functionally related gene groups (BINs), as they are defined by the MapMan ontology^29^. Important immunity-related or regulatory BINs, enriched in up- or down-regulated genes of PVY-vs. mock-inoculated plants for all three genotypes, are listed in Table 1. Up-regulated genes enriched uniquely in TGA2.1-NahG included isoprenoid metabolism-related genes, PHD finger and PHOR1 transcription factors, and peroxidases (Table 1). On the other hand, the C2C2(ZN) DOF transcription factors were enriched in down-regulated genes in TGA2.1-NahG plants. PVY-regulation of cytokinin and jasmonate metabolism was lost in TGA2.1-NahG compared with the other two genotypes, as was the up-regulated expression of WRKY transcription factors (Table 1).

**Table 1.** Selected functional groups (BINs) enriched in up- or down-regulated genes in TGA2.1-NahG, NahG and NT plants after PVY infection. FDR corrected q-value < 0.05. (+), enriched in up-regulated genes; (−), enriched in down regulated genes.

| BIN | Functional group | TGA2.1-NahG | NahG | NT |
| --- | --- | --- | --- | --- |
| Secondary metabolism |  |  |  |  |
| 16.1.2 | mevalonate pathway | + |  |  |
| 16.1.5 | terpenoids | + |  |  |
| 16.10 | simple phenols |  |  | + |
| Hormone metabolism |  |  |  |  |
| 17.1.3 | abscisic acid-regulated | – | – |  |
| 17.2 | auxin | – | – |  |
| 17.2.3 | auxin-regulated | – | – |  |
| 17.4.1 | cytokinin |  | – | – |
| 17.7 | jasmonate |  | + | + |
| 17.8 | salicylic acid |  |  | + |
| Stress – biotic |  |  |  |  |
| 20.1.7 | PR proteins | + | + | + |
| 20.1.7.1 | PR-1 | + |  |  |
| 20.1.7.3 | PR-3/PR-4/PR-8/PR-11 | + | + | + |
| Miscellaneous |  |  |  |  |
| 26.12 | peroxidases | + |  |  |
| 26.21 | protease inhibitor/seed storage/lipid transfer proteins |  | – |  |
| Transcription regulation |  |  |  |  |
| 27.3.32 | WRKY transcription factors |  | + | + |
| 27.3.63 | PHD finger transcription factors | + |  |  |
| 27.3.64 | PHOR1 transcription factors | + |  |  |
| 27.3.8 | C2C2(Zn) DOF transcription factors | – |  |  |
| DNA synthesis – chromatin structure |  |  |  |  |
| 28.1.3 | histone | + | + | + |
| 28.1.3.2 | histone core | + | + | + |
| 28.1.3.2.1 | histone core H2A proteins | + | + |  |
| 28.1.3.2.3 | histone core H3 proteins | + | + | + |
| Protein degradation |  |  |  |  |
| 29.5.11.20 | ubiquitin-proteasome | + | + |  |
| Signaling – receptor kinases |  |  |  |  |
| 30.2.8.1 | leucine-rich repeat VIII (type 1) |  |  | – |
| 30.2.16 | *Catharanthus roseus*-like RLK1 |  |  | + |
| 30.2.17 | DUF26 |  |  | + |
| 30.2.19 | legume-lectin | + |  | + |
| 30.2.99 | miscellaneous | + |  | + |

### StTGA2.1 and StTGA2.3 activate the class III peroxidase StPRX07

As the expression of several peroxidases was up-regulated after *StTGA2.1* overexpression in PVY-infected plants (Table 1, Supplementary Table 4), we recognized them as potential direct targets of StTGA2.1. To test our hypothesis, we selected three class III peroxidases (StPRX, Supplementary Table 5), *StPRX07* (*Sotub09g020950*), *StPRX15* (*Sotub02g035680*), and *StPRX46* (*Sotub03g007840*), which were up-regulated in TGA2.1-NahG compared with NahG (GEO accession GSE196078, Supplementary Table 6), for further analysis. Analysis of their promoter regions revealed predicted TGA-binding motifs between 450 and 750 bp upstream of the transcription start site (Fig. 4a).

**Fig. 4.**
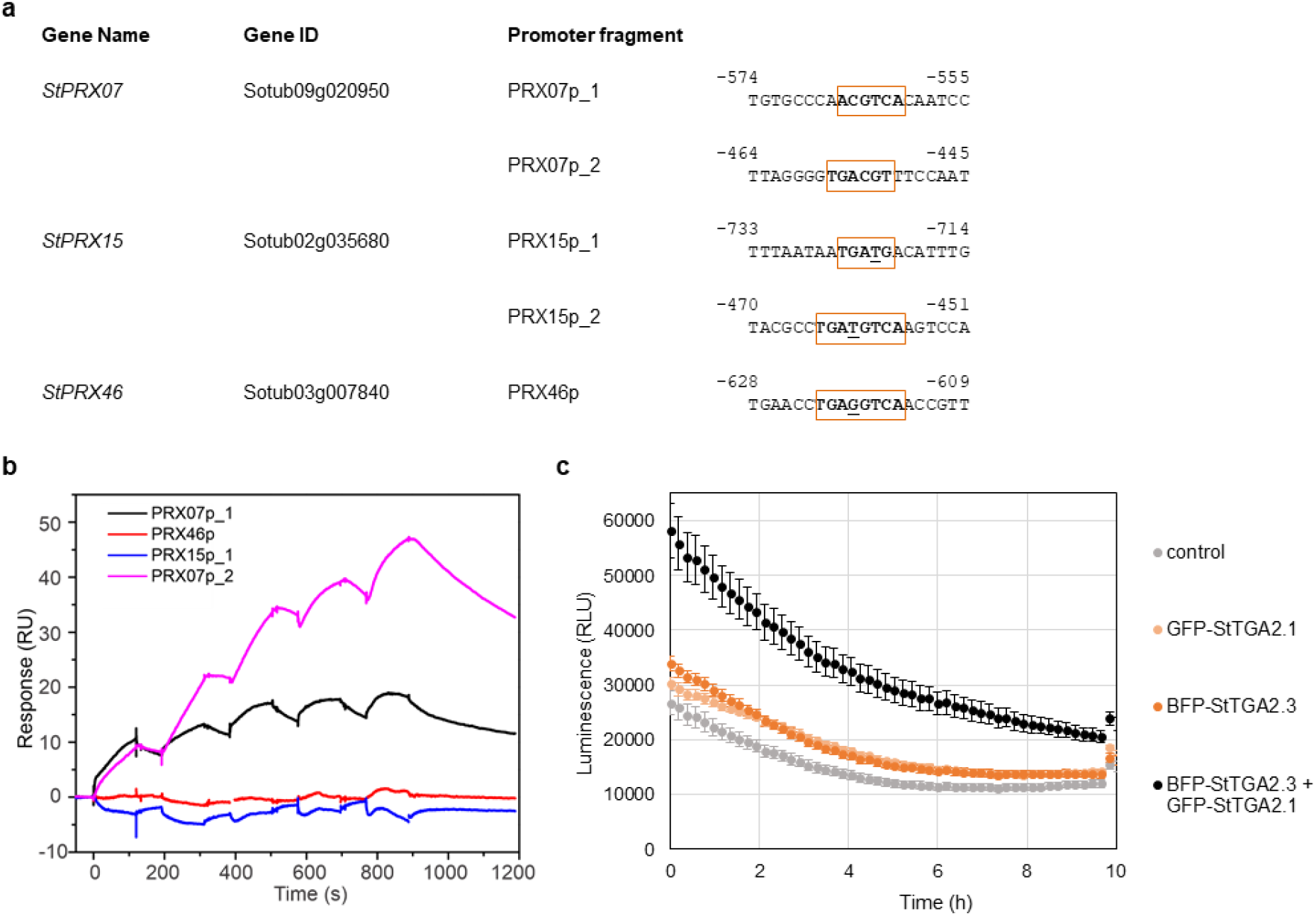
StTGA2.1, together with its interactor StTGA2.3, activates expression of StPRX07. **a,** TGA-binding motifs in selected *StPRX* promoters. The predicted motifs are boxed and the nucleotides, differing from the core TGACG(T) sequence, its reverse complement or the TGACGTCA palindrome, are underlined. Numbers indicate position upstream of transcription start site. **b,** Surface plasmon resonance results, showing the interaction between StTGA2.3 and chip-immobilized PRX15p_1, PRX46p, PRX07p_1 or PRX07p_2 DNA fragments, bound to the chip at ∼38, 41, 65, or 53 response units (RU), respectively. Representative sensorgrams are shown. **c,** Transactivation assay results, showing *in planta StPRX07* promoter activation by GFP-tagged StTGA2.1 (light orange), BFP-tagged StTGA2.3 (dark orange) or a combination of both (black). BFP or GFP-tagged controls and their combination (control) were used to detect the basal promoter activity (grey). Average values ± standard error of 18 biological replicates in the first 10 h of measurement are shown. The experiment was repeated twice with similar results (Supplementary Fig. 8b).

To investigate the ability of StTGAs to bind these motifs, we first tested whether the StTGA2.3 and StTGA2.1 proteins could bind to four candidate DNA fragments from *StPRX* promoter regions, PRX07p_1, PRX07p_2, PRX15p_1 and PRX46p (Fig. 4a), using surface plasmon resonance. Titration of a recombinant StTGA2.3 over chip-immobilized PRX07p_1 and PRX07p_2 fragments, carrying the predicted TGA-binding motifs of the *StPRX07* promoter, resulted in a dose-dependent increase in response, compared to reference (Fig. 4b). Interaction with PRX15p_1 and PRX46p fragments was negligible (Fig. 4b). As predicted by the absence of the basic region, we did not measure any interaction between the His_6_-tagged StTGA2.1 and the tested DNA (Supplementary Fig. 8a). These results suggest that StTGA2.3, but not StTGA2.1, binds specifically to the TGA-binding motifs in the *StPRX07* promoter. Finally, we tested the ability of StTGA2.3 and StTGA2.1 to activate the *StPRX07* promoter *in planta*, using a transient transactivation assay^30^. For this purpose, the 2.95 kbp long promoter region upstream of the *StPRX07* start codon, containing both predicted TGA-binding motifs, was fused to a luciferase (*LucF*) coding sequence. GFP-tagged StTGA2.1 and BFP-tagged StTGA2.3 were then co-expressed with the reporter construct, confirmed by confocal microscopy. Co-expression of the reporter construct with StTGA2.3 induced the expression of *StPRX07*::*LucF* by approximately 20% compared with basal promoter activity, whereas co-expression with StTGA2.1 resulted in only minor induction (Fig. 4c, Supplementary Fig. 8b). In contrast, more than two-fold induction in promoter activity was observed when co-expressed with both StTGA2.1 and StTGA2.3 (Fig. 4c, Supplementary Fig. 8b). These results indicate that strong activation of *StPRX07* promoter is achieved only when both StTGA2.3 and StTGA2.1 are present.

### StTGA2.1 N-terminus likely contributes to protein interactions and alters TGA binding to DNA

Comparative structural analysis using AlphaFold (AF)^31^ revealed important singularities in the molecular architecture of StTGA2.1, mostly contained in its N-terminus. In the AF models of StTGA2.2 and StTGA2.3, the intrinsically disordered N-terminus is followed by an α- helical bZIP domain, which includes several basic residues and three heptads that comprise the leucine zipper (Fig. 5a,b, Supplementary Fig. 9). In contrast, StTGA2.1 has a short N-terminus with low helical propensity due to the Pro25 α-helix breaker, harbouring only the basic residues Arg20, Arg30, and Arg24, and lacking the first and most of the second leucine zipper heptad (Fig. 5a,b). The conserved hydrophobic residues Leu34, Val31, and Phe38 may contribute to protein dimerization by forming a partial zipper that could be stabilized by the hydrophobic Val28 and Val29. Persistent contacts identified in molecular dynamics (MD) simulations of StTGA2.2 and StTGA2.3 homo- and heterodimers are preserved in StTGA2.1 and involve the said hydrophobic residues (Fig. 5b, Supplementary Fig. 10).

**Fig. 5.**
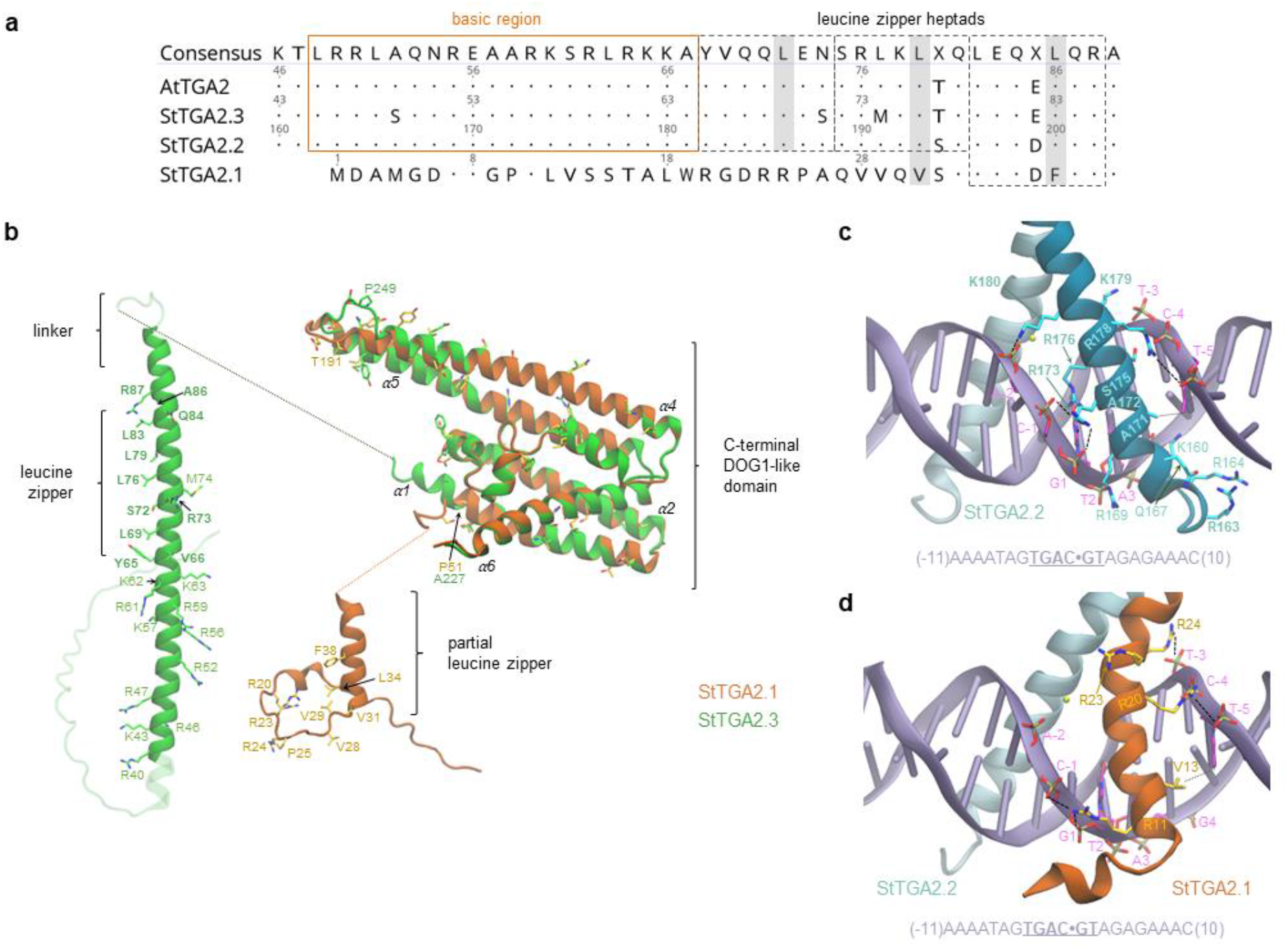
Comparative structural analysis and simulations of StTGA2.1 N-terminus interactions with StTGA2.2 and StTGA2.3 bZIP domains. **a**, Protein sequence alignment of StTGA2.1 N-terminus with AtTGA2, StTGA2.2 and StTGA2.3 bZIP domains. The basic region (orange box) and the leucine zipper heptads (grey dashed boxes) are indicated. Conserved amino acids, in respect to the consensus sequence, are marked with dots. StTGA2.1 contains hydrophobic residues (Val31 and Phe38) in two out of three Leu positions in the heptads (grey) and has a completely conserved third heptad. **b**, Molecular architectures of StTGA2.1 (orange) and StTGA2.3 (green). The StTGA2.3 bZIP domain (aa 40-95), and StTGA2.1 N-terminus (aa 1-45) are shown. The C-terminal region is highly conserved between StTGA2.1 (aa 47-240), StTGA2.2 (aa 222-446), and StTGA2.3 (aa 105-327). Amino acid residues that are discussed in this study are represented as liquorice and labelled. Those forming persistent contacts in the leucine zipper, according to molecular dynamics (MD) simulations, are shown in bold. Basic amino acid residues that may contribute to DNA-binding are depicted and labelled. Fully non-conservative substitution sites in the putative DOG1 domains are also represented as liquorice. **c,** Representative snapshot of the MD simulations of the DNA-bound StTGA2.2 homodimer and **d,** StTGA2.2-StTGA2.1 heterodimer. The DNA double helix is represented in violet. The DNA sequence is shown at the bottom and the binding site core is underlined. A dot is used as a reference at the centre of the sequence for numbering the nucleotide residues. StTGA2.1 (orange) and StTGA2.2 (cyan) are represented as cartoon. Salt bridges and hydrogen bonds between protein and DNA are indicated with black dashed lines. Hydrophobic contacts are indicated as dotted lines. Amino acid residues forming persistent interactions are labelled in bold. The presence of a hexacoordinated Mg^2+^ was assumed (yellow sphere), based on its importance for CREB- bZIP^32^.

We then inquired if the StTGA2.1 N-terminus binds to DNA. Based on prior knowledge of AtTGA2 and its cognate TGACG motif^10^, we modelled the DNA-bound StTGA2.2- StTGA2.2 and StTGA2.2-StTGA2.1 dimers and, via MD simulations, identified the key DNA-binding residues (Fig. 5c,d, Supplementary Tables 7,8). Persistent interactions in the StTGA2.2 homodimer mostly correspond to salt bridges, formed between the bZIP basic residues and the DNA phosphate groups, and do not explain the StTGA2.2 motif-binding specificity. In contrast, sequence specificity is provided by hydrophobic contacts between the StTGA2.2 Ala172 residue and the DNA T2 methyl group or between Ala171 and T-4 or T-5. Indeed, most of the predicted TGA-binding motifs (Fig. 4a) have a thymine or adenine in these positions. While the contacts involving StTGA2.2 in the StTGA2.1-StTGA2.2 heterodimer are highly preserved, the StTGA2.1-DNA interactions are dramatically reduced, with only Arg11, Arg20, and Arg24 forming persistent contacts with the DNA phosphate groups (Fig. 5d, Supplementary Table 8). Moreover, the StTGA2.1 partial zipper positions the last basic residue (Arg24) more distant to the DNA compared to StTGA2.2 (Lys180), breaking the protein-DNA complex symmetry.

Another important singularity of StTGA2.1 is that its partial zipper connects directly to the putative DOG1 domain, while the two domains are connected by a 13 aa peptide linker in StTGA2.2 and StTGA2.3. This may greatly influence its interdomain conformational flexibility. Pro51 in StTGA2.1 (Ala227 and Ala110, in StTGA2.2 and StTGA2.3, respectively), disrupts the DOG1 α-helix 1 (α1) and forms a shorter, disordered linker that could somehow compensate for this absence. These results indicate that the compact molecular architecture of StTGA2.1, which causes an asymmetric distribution of basic residues in the StTGA2.1-StTGA2.2 heterodimer, significantly distorts the DNA conformation near the binding site, supporting strong promoter activation upon binding of the heterodimer compared with binding of the homodimer.

## DISCUSSION

TGAs are involved in modulation of various cellular processes, acting as positive or negative regulators of gene expression^3^. Their structural features provide the basis for their functional variability, defining their subcellular localization, target recognition, DNA-binding, as well as their ability to form dimers, oligomers or interact with other proteins (*under review, Š. T., K.G. and A.C.*). In Arabidopsis, all ten AtTGAs share a highly conserved bZIP domain, essential for establishing specific interactions with DNA. Here, we report on the structural-functional relationship of a mini-TGA from potato, StTGA2.1, which lacks a full bZIP but still acts at target gene activation. Mini-TGAs were identified in potato (this study), tomato^24,25^, and strawberry^19^, but are not found in Arabidopsis. The potato and tomato mini- TGAs are orthologous to AtTGA clade II members, although they form a separate phylogenetic group together with StTGA2.2 and SlTGA2.2 (Fig. 1a), suggestive of an earlier diversification in evolutionary history.

Homo- and heterodimerization of StTGA2.1 (Fig. 3a,b, Supplementary Fig. 3), which contains only a part of an already short and presumably unstable TGA zipper region^33^, corroborates the findings of Boyle *et al.* (2009), who established that the leucine zipper is not crucial for dimerization of AtTGA2^10^. Instead, StTGA2.1 dimerization is likely mediated by interactions involving its C-terminal region, previously reported to contain a dimer stabilization region^34^. StTGA2.1 nuclear localization allows its role in gene regulation, however its unusually broad localization pattern (Fig. 3c, Supplementary Fig. 5) suggests that it might perform different tasks, as has been shown for other plant transcription factors with intrinsically disordered regions^35^. TGAs also interact with different proteins present in both nuclei and cytoplasm, including NPR cofactors^6,10^, which was already confirmed for StTGA2.1 (Supplementary Fig. 4), glutaredoxins^36^, and calmodulins^37^. Thus, StTGA2.1 could modulate the function of these partners by sequestering them into inactive complexes, as was proposed for AtTGA2 and NPR1^38^. Multiple functions of StTGA2.1 are supported also by the diversity of detected transcriptional changes following exposure to pathogen infection (Table 1, Supplementary Table 4, GEO accession GSE196078).

Multiple studies have evaluated the influence of clade II TGA dominant-negative bZIP mutants on plant immunity during bacterial infection, leading to contradictory results^38–40^. Reactive oxygen species (ROS) act as signalling molecules in biotic stress^41^. Early ROS production is central to plant defence and TGAs have previously been associated with cellular redox control, physically interacting with or regulating the expression of CC-type glutaredoxins^25,36,42^. Furthermore, clade II TGAs modulate the expression of glutathione-S-transferases in ROS-processing responses to UV-B stress^43^, while clade IV AtTGAs are regulated by flg22-induced ROS production^44^. Here we show that the synergistic activity of TGAs regulates the expression of yet another group of enzymes involved in ROS-metabolism, the class III peroxidases (Fig. 4b,c, Supplementary Fig. 8). Class III peroxidases are haem-containing glycoproteins, secreted to the apoplast or localized in vacuoles^45^. Among them, AtPRX33 and AtPRX34 proved vital for apoplastic ROS production in response to flg22 and elf26^46^. Most of the StPRX protein sequences from the peroxidase functional group^29^ contain predicted secretory signal peptides (Supplementary Table 5), indicating StTGA2.1 could affect apoplastic ROS production in plant defence.

Transcription factor cooperativity is essential in eukaryotic transcription regulation and can arise through various mechanisms, involving protein-protein and/or protein-DNA interactions^47^. Previous studies have shown that TGA mutants, impaired in DNA-binding through diverse modifications of the bZIP domain, prevent DNA-binding of wild type homologues^39,40,48^, which somewhat opposes the cooperative activation via an StTGA2.1- StTGA2.3 complex. Compared to homodimers of its paralogues, our molecular dynamics simulations suggest that the asymmetrical distribution of basic residues in the bZIP-like domain in the StTGA2.1-StTGA2.2 heterodimer significantly distorts the DNA conformation near its binding site (Fig. 5). We hypothesize that StTGA2.1 dramatically affects the overall conformation of the regulatory complex due to its compact molecular architecture.

In conclusion, we show that, although mini-TGAs are not able to bind DNA on their own, their unusual structure supports diverse functionalities, such as allowing the induction of class III peroxidases in immune signalling. We thus provide evidence that truncation in evolution of genes does not necessary lead to a loss-of-function phenotype. Instead, additional functions can be attained. Through this, we shed additional perspective on immune signalling in non- model species, as Arabidopsis does not encode such proteins.

## MATERIALS AND METHODS

### *In silico* sequence and structural analysis

TGA transcription factor orthologues from potato were identified based on orthologue information included in the GoMapMan database^29^. The initial list was further pruned based on protein sequence alignments created with Geneious Alignment in Geneious Prime 2020.1.1 (https://www.geneious.com) and BLAST results to exclude technical errors of orthologue detection and sequencing. Identified StTGAs are listed in Supplementary Table 1. Basic protein information was calculated using the ExPASy ProtParam tool^49^. Protein domain prediction was performed with ExPASy Prosite^50^. Protein sequences of SlTGAs ^24,25^ were retrieved from the Sol Genomics Network^51^ and sequences of AtTGAs from The Arabidopsis Information Resource^52^.

For the phylogenetic analysis, the sequences were aligned with MAFFT^53^, using the L-INS-I iterative refinement method, and the alignment used for a maximum-likelihood phylogenetic tree construction in MEGA-X^54^, using the Jones-Taylor-Thorton matrix-based model^55^ and 1,000 bootstrap repetitions. The rice OsTGA2.1^26^ protein sequence (Q7X993) was retrieved from UniProtKB (https://www.uniprot.org/) and used as tree root. For sequence similarity visualization, the protein sequences were aligned with Geneious Alignment in Geneious Prime 2020.1.1 (https://www.geneious.com). Potato peroxidases were identified with protein sequence BLAST against the RedoxiBase database^56^ and the secretory signal peptides were predicted with SignalP 5.0^57^ (Supplementary Table 5). Predictions of transcription factor binding motifs in promoter sequences were performed with TRANSFAC^58^ and predictions of transcription start sites with TSSFinder^59^.

Structural models of StTGA2.1, StTGA2.2, and StTGA2.3 were generated with AlphaFold^31^. The top-ranked models were selected. The VMD (Visual Molecular Dynamics, version 1.9.4a48) molecular visualization program was used for visual analysis and structural alignment of protein models.

### Molecular dynamics simulations

The initial homo- and heterodimeric configurations of StTGA2.1-StTGA2.2, StTGA2.2- StTGA2.2 and StTGA2.2-StTGA2.3 N-terminal fragments were defined using the crystal structure of CREB bZIP-CRE (PDB id: 1DH3) as template^60^. Corresponding amino acid changes to the template were done using the VMD psfgen plugin, preserving the coordinates of the backbone and C_β_ atoms. StTGA2.1, StTGA2.2 and StTGA2.3, are truncated, keeping the amino acids 1-43, 159-206, and 42-89, respectively. The N-termini of StTGA2.2 and StTGA2.3 are capped with N-methylamide and the C-termini of the three proteins with acetyl. For simulations of DNA-bound StTGA2.1-StTGA2.2 and StTGA2.2-StTGA2.2, the DNA fragment from the template crystal structure was kept and the nucleotides were modified using psfgen. The final DNA sequence corresponds to the TGACGT motif, complementary to the linker scan 5 element and its adjacent regions of Arabidopsis *PR-1* promoter *as-1*-like sequence^61^ (Fig. 5c,d). The crystal Mg^2+^ cation and the six coordinated water molecules were kept.

GROMACS-2020^62^ was used to prepare inputs and run molecular dynamics (MD) simulations. The simulation boxes were generated as an octahedron, defining a solvation layer of 10 Å minimum thickness around the molecular complex. 0.15 M NaCl was used to establish electroneutrality. Protonation states were defined for pH 7.0. Amber ff99SB^63^ and ff14SB9^64^ were used to describe the protein in the free and DNA-bound TGA dimers, respectively, and PARMBSC1^65^ was used to describe the DNA. TIP4P-D^66^ or SPC^67^ was used to describe water molecules in the simulations of the free and DNA-bound TGA dimers, respectively. CHARMM-formatted topology and parameter files were converted to GROMACS input files using the VMD plugin TopoGromacs^68^.

The MD simulations were performed on the Summit supercomputer at the Oak Ridge Leadership Computing Facility. Energy minimization was performed for all systems with steepest descent. Periodic electrostatic interactions were treated with the particle mesh Ewald method^69^. LINCS^70^ was used to constrain bonds involving hydrogen atoms.

Similar protocols of simulation were applied for the free and DNA-bound TGA dimers. Preceding the classical simulations, we performed long equilibration runs of 315 ns as part of our protocol adapted from the MD simulation-based method of structural refinement described by Heo *et al.* (2021)^71^. In this protocol, potential sampling is accelerated with hydrogen mass repartitioning and by applying fairly high temperatures. Weak position restraint potentials were applied for minimum bias and to compensate for the high thermal energy. Velocity Langevin dynamics was performed using a friction constant of 1 ps^-1^. During the equilibration phase, position restraints applied to C_α_ atoms in the leucine heptads were gradually released and the temperature gradually increased, reaching the maximum of 360 K (Supplementary Table 9). After long sampling at 340 K and 320 K, a final phase of equilibration is conducted at 298.15 K, the temperature of the following production runs. During the final equilibration phase, flat-bottom harmonic restraint potentials were applied, using a force constant of 0.25 kcal/mol/Å^2^ and a flat-bottom width of 4 Å. To adjust box size, part of the equilibration phase was conducted in the *NpT* ensemble, using the Berendsen barostat^72^ applying a compressibility of 4.5 x 10^-5^ bar^-1^ and a time constant of 1.0 ps. In the final phase of equilibration, the atomic velocities were assigned from a Maxwell-Boltzmann distribution using random numbers of seed. The production runs of free and DNA-bound dimers consisted of five unbiased independent simulations of 128 ns and 200 ns, respectively. The position restraint potential applied to Mg^2+^ and its coordinated water molecules was kept during these simulations.

For simulation analysis, the VMD plugin Hbonds was used to compute hydrogen bond statistics. The geometric criteria adopted are a cut-off of 3.0 Å for donor-acceptor distance and 20 ° for acceptor-donor-H angle. In Fig. 5c,d, salt bridges and hydrogen bonds between protein DNA occurring during more than 10% of the simulation time are shown. Persistent contacts were identified using the VMD plugin Timeline. In Fig. 5c,d, amino acid residues involved interactions or hydrophobic contacts persisting for more than 30% of the simulation time are shown. Grace was used for plots (https://plasma-gate.weizmann.ac.il/Grace/).

### Plant material

Potato (*Solanum tuberosum* L.) non-transgenic cultivar Rywal (NT) and Rywal-NahG (NahG), a transgenic line impaired in SA accumulation due to salicylate hydroxylase expression^23^, were used in this study. Plants were propagated from stem node tissue cultures and transferred to soil two weeks after node segmentation, where they were kept in growth chambers under controlled environmental conditions at 22/20 °C with a long-day (16 h) photoperiod of light (light intensity 4000 lm/m^2^) and 60-70% relative humidity. Tobacco *Nicotiana benthamiana* plants were grown from seeds and kept in growth chambers under the same conditions.

### DNA constructs

Full-length coding sequences (cds) of StTGA2.1 (Genbank accession number OM569617), StTGA2.2 (Genbank accession number OM569618), StTGA2.3 (Genbank accession number OM569619), StNPR1 (Genbank accession number OM569620) and StNPR3/4 (Genbank accession number OM569621) were amplified from potato cultivar Rywal cDNA and inserted into the pJET1.2/blunt cloning vector using the CloneJET PCR Cloning Kit (Thermo Scientific, USA), following the manufacturer’s instructions.

The selected genes were subsequently cloned into pENTR D-TOPO vector using pENTR™ Directional TOPO® Cloning Kit (Invitrogen, USA) and recombined through LR reaction using the Gateway® LR Clonase TM II Enzyme Mix (Invitrogen, USA) into several Gateway destination vectors (VIB, Belgium). For co-immunoprecipitation experiments, localization studies and transactivation assays, the StTGA2.1, StTGA2.2 and StTGA2.3 cds were inserted into pH7FWG2 and pJCV52 expression vectors^73^ to produce proteins with C-terminal enhanced green fluorescent protein (GFP) and hemagglutinin A (HA) fusions, respectively. For transactivation assays, StTGA2.3 was fused with a C-terminal mTagBFP2 (from Addgene plasmid # 102638)^74^ blue fluorescent protein (BFP) prior to cloning into pENTR D-TOPO vector (Invitrogen, USA) and subsequently recombined into the pK7WG2 vector^73^. A short linker of six Gly residues was introduced between the StTGA2.3 and BFP sequence. BFP fused with an N7 nuclear localization signal^75^ (N7-BFP) was recombined into pK7WG2^73^ as control.

For overexpression experiments, the StTGA2.1 cds was amplified with primers harbouring *XhoI* and *Spe*I restriction enzyme cleavage sites and inserted into the pTA7002 vector^28^, enabling glucocorticoid-inducible gene expression *in planta*, through restriction-ligation cloning.

For the yeast two-hybrid assays, the cds of StTGA2.1, StTGA2.2, StTGA2.3, StNPR1 and StNPR374 were amplified and inserted into the pGBKT7 (bait) yeast expression vector through *in vivo* cloning with Matchmaker Gold Yeast Two-Hybrid System (Clontech, USA), to produce proteins with an N-terminal Gal4 DNA-binding domain. StTGA2.1, StNPR1 and StNPR3/4 were inserted also into the pGADT7 (prey) vector (Clontech, USA), to produce proteins with an N-terminal Gal4 activation domain, using the same cloning system.

Promoter sequences of *StPRX07* (Genbank accession number OM569622), *StPRX15* (Genbank accession number OM569623) and *StPRX46* (Genbank accession number OM569624) were amplified from potato cultivar Rywal genomic DNA and inserted into the pENTR D-TOPO vector (Invitrogen, USA). The *StPRX07* promoter sequence was subsequently recombined through LR reaction into the pGWB435 Gateway vector^76^, as described above, inserting the promoter upstream of a luciferase reporter (*LucF*).

For recombinant protein production, the StTGA2.1 cds was inserted into the pMCSG7 bacterial expression vector^77^ by ligation-independent cloning^78^ to produce a protein with an N-terminal hexahistidine (His_6_) tag. The cds of StTGA2.3 was amplified using primers, enabling the digestion-ligation reaction with the *Bsa*I restriction enzyme. Three silent mutations were introduced into its sequence, to remove two native *Bsa*I restriction sites. The amplified fragment was subsequently ligated into the pEPQD0KN0025 acceptor backbone (Addgene plasmid #162283)^79^, together with pEPQD0CM0030 (Addgene plasmid #162312)^79^, which adds an additional GS peptide to the protein C-terminus.

All primer pairs used in the cloning procedure are listed in Supplementary Table 10. Sequence verification was performed with Sanger sequencing (Eurofins Genomics, Germany).

### Transient expression assays

Homemade electrocompetent *Agrobacterium tumefaciens* GV3101 cells were transformed with prepared constructs by electroporation. Transformants were used for agroinfiltration of the bottom three fully developed leaves of 3-4 week old *N. benthamiana* plants, as described previously^80^. In cases of co-transformation with agrobacteria carrying different constructs, the 1:1 ratio was applied. An equal volume of agrobacteria carrying *p19* silencing suppressor (kindly provided by prof. Jacek Hennig, PAS, Poland) was added to the mixture. Agrobacteria carrying *p19* only were used as controls.

### Confocal microscopy

Protein fluorescence was visualized three to five days after transient *N. benthamiana* transformation. For protein localization, the Leica TCS SP5 laser scanning confocal microscope mounted on a Leica DMI 6000 CS inverted microscope with an HC PL FLUOTAR 10x objective or HCX PL APO lambda blue 63.0×1.40 oil-immersion objective (Leica Microsystems, Germany), using the settings described previously^81^. The red Histone 2B-mRFP1 (H2B-RFP) nuclear marker^82^ was used to visualize cell nuclei. For co-immunoprecipitation and transactivation assays, the protein fluorescence was confirmed with the Leica TCS LSI macroscope with Plan APO 5x and 20x objectives (Leica Microsystems, Germany), using the settings described previously^83^. The green, blue or red fluorescent protein fluorescence was excited using 488 nm, 405 nm and 543 nm laser lines, respectively. The emission was measured in the window of 505-520 nm for GFP, 450-465 nm for BFP, 570-630 nm for H2B-RFP and 690-750 nm for autofluorescence. The Leica LAS AF Lite software (Leica Microsystems, Germany) was used for image processing.

### Yeast two-hybrid assay

Bait (containing *StTGA2.1*, *StTGA2.2*, *StTGA2.3*, *StNPR1* or *StNPR3/4* cds), and prey (containing *StTGA2.1*, *StNPR1* or *StNPR3/4* cds) construct combinations were transformed into the Y2H Gold strain using the Matchmaker Gold Yeast Two-Hybrid System (Clontech, USA) and the transformants selected on control SD media without Leu and Trp (-L-W). Interactions were analysed on selection SD media without Leu, Trp, His and adenine, with added X-α-Gal and Aureobasidin A (-L-W-H-A+Xgal+Aur). The proteins were tested for autoactivation through co-transformation of bait constructs with an empty prey vector. To evaluate the strength of interaction, saturated yeast culture dilutions (10^-1^, 10^-2^ and 10^-3^) were spotted onto selection media. To evaluate the effect of SA on the strength of interaction, the dilutions were spotted onto selection media containing 0.1 mM or 1.0 mM SA.

### Co-immunoprecipitation assay

HA or GFP-tagged StTGA2.1, StTGA2.2 and StTGA2.3 were transiently expressed in *N. benthamiana* leaves in different combinations. The fluorescence of GFP-tagged proteins was confirmed with confocal microscopy after 4 days. Total proteins were extracted from ∼500 mg leaf material with immunoprecipitation (IP) buffer, containing 10 mM Tris-HCl, pH 7.5, 150 mM NaCl, 2 mM MgCl_2_, 1 mM dithiothreitol and 1x EDTA-free Protease Inhibitor Cocktail (Roche, Switzerland), followed by 1 h incubation with GFP-Trap® Magnetic Agarose beads (ChromoTek, Germany) at 4 °C. The beads were washed three times with IP buffer and eluted into SDS-PAGE loading buffer, containing 100 mM Tris-HCl, pH 6.8, 4% (w/v) SDS, 0.2% bromophenol blue, 20% (v/v) glycerol and 200 mM dithiothreitol. The immunoprecipitated proteins and protein extracts were analysed by SDS-PAGE and Western blot, using anti-GFP (diluted 1:3.000 or 1: 5.000, Invitrogen, USA) and anti-HA (diluted 1:1.000, ChromoTek, Germany) antibodies.

### Generation of StTGA2.1 overexpression plants

Transgenic NahG-TGA2.1 plants were obtained by stable transformation of the Rywal-NahG potato genotype^23^. Electrocompetent *A. tumefaciens* strain LBA4404 was electroporated with the pTA7002 vector^28^ carrying the *StTGA2.1* cds, as described above. Agrobacteria were used for stable transformation of sterile plantlet stem internodes from tissue culture, as described previously^84^. Plantlets grown on regeneration media plates with hygromycin selection were sub-cultured in order to generate independent transgenic lines. Transgenic lines were confirmed with PCR (Supplementary Table 10). Lines 7 and 12 were selected for further analysis.

### Virus inoculation and plant treatments

Three to four weeks old potato plants were inoculated with GFP-tagged infectious PVY clones PVY^N605^–GFP^85^ or PVY^N605^(123)-GFP^84^ or mock inoculum, as described previously^86^. To induce gene overexpression, plants were treated with dexamethasone (DEX) foliar spray solution containing 30 μM DEX and 0.01 % (v/v) Tween-20 or control spray solution without DEX (control), 3 h prior to virus inoculation, 3 h after virus inoculation and every day post inoculation until sampling.

### Gene expression analysis with qPCR

For gene expression analysis, total RNA isolation, reverse transcription and qPCR were performed as described previously^23^. DEX-induced *StTGA2.1* overexpression in fully developed leaves of TGA2.1-NahG transgenic lines was confirmed 3 h after DEX treatment using a qPCR assay targeting *StTGA2.1* cds. The leaves of three DEX-treated plants and two or three non-treated plants were sampled, one leaf per plant. For PVY abundance analysis, PVY-infected leaves of DEX-treated TGA2.1-NahG, NahG and NT genotypes were sampled at 3, 5 and 7 dpi or 3, 7 and 10 dpi. For each genotype and treatment, three plants were analysed, sampling one leaf per plant per dpi. PVY abundance and *StTGA2.1* expression were quantified using two sample dilutions and a relative standard curve method by normalization to the endogenous control *StCOX1* with quantGenius (http://quantgenius.nib.si)^87^. A two- tailed *t*-test was used to compare treatments, when applicable. The qPCR analysis was performed for both TGA2.1-NahG transgenic lines.

RNA sequencing results were validated technically and biologically with qPCR, as described above. For technical validation, the expression of *StACX3*, *StCS*, *StPti5, StPRX28* and *StTGA2.1* was followed. Biological validation was performed in an independent experiment repetition with both TGA2.1-NahG transgenic lines and following gene expression of *StPRX07*, *StPRX15*, *StPRX46*, *StTGA2.1* and *PVY*. *StCOX1* and *StEF-1* were used for normalization in both cases, as described above. A two-tailed *t*-test was used to compare treatments, when applicable.

All primers and probes used for qPCR analysis together with the target gene IDs are listed in Supplementary Table 11. New qPCR assays, targeting *StPRX07*, *StPRX15*, *StPRX46*, *StTGA2.1* and *StPti5* were designed with Primer Express v2.0 (Applied Biosystems, USA), using the sequences from the potato reference genome^88^, cultivar Rywal cds, and cultivar Rywal and cultivar Désirée reference transcriptomes^89^.

### RNA sequencing analysis

For RNA sequencing, 2-25 early visible lesions and their immediate surroundings were sampled from PVY-inoculated leaves of DEX-treated TGA2.1-NahG, NahG and NT plants and control-treated TGA2.1-NahG plants at 4 dpi, as described previously^27^. About 20-30 sections of comparable size were sampled from mock-inoculated leaves as controls. Three plants per genotype per treatment were analysed, pooling together all lesions or mock- sections from one leaf per plant. Total RNA was isolated as described previously^27^. Strand-specific library preparation and sequencing were performed by Novogene (HongKong), using the NovaSeq platform (Illumina) to generate 150-bp paired-end reads. Read quality control was performed using FastQC^90^. The presence of contaminant organism reads was determined using Centrifuge^91^. Reads were mapped to the reference group Phureja DM1-3 potato genome v4.04^88^ using the merged PGSC and ITAG genome annotation^89^ and counted using STAR^92^ with default parameters. Differential expression analysis was performed in R using the limma package^93^. Raw and normalized read counts as well as a processed data table were deposited at GEO under accession number GSE196078. Genes with Benjamini-Hochberg FDR adjusted p-values < 0.05 and |log_2_FC| ≤ −1 were considered statistically significantly differentially expressed. The Venn diagram was drawn, according to results obtained with the Gene List Venn Diagram tool (http://genevenn.sourceforge.net/).

Gene Set Enrichment Analysis^94^ was performed using non-filtered normalized counts to search for regulated processes and functionally related gene groups, altered significantly by virus inoculation in different genotypes (FDR corrected q-value < 0.05) using MapMan ontology^29^ as the source of gene groups.

### Targeted genomic sequencing

Genomic DNA was isolated from potato cultivar Rywal leaves using the DNeasy Plant Mini Kit (Qiagen, Germany). Two sets of primers were designed to target the region of interest (Supplementary Fig. 1a, Supplementary Table 12). Droplet-based PCR-free target region enrichment, library preparation using the SQK-LSK109 kit (Oxford Nanopore Technologies, United Kingdom) and long-read sequencing on the MinION platform using the R9.4.1-type flow cell was performed by Samplix (Denmark). Nanopore read basecalling was performed using Guppy 4.2.2. The reads were error corrected with NECAT^95^ setting GENOME_SIZE=100000000 and PREP_OUTPUT_COVERAGE=20000. Chimeric reads were split using Pacasus^96^ and all reads designated as “passed” were mapped to the group Phureja DM1-3 potato genome v6.1^97^ using Minimap2^98^. The obtained BAM file was indexed and sorted using SAMtools^99^. Raw Nanopore reads were deposited at SRA under accession number PRJNA803339.

### Transactivation assay

GFP-tagged StTGA2.1, BFP-tagged StTGA2.3 and their combination were transiently expressed in *N. benthamiana* leaves, with N7-BFP and either a GFP-tagged SNF-related serine/threonine-protein kinase (StSAPK8) or an empty pH7FWG2 vector^73^ as controls. Protein fluorescence was confirmed with confocal microscopy after 3-5 days. The transactivation assays were performed as described previously^30^. In brief, 0.5-cm-diameter leaf discs were sampled at 4 dpi and pre-incubated in MS liquid media with 35 μM D- luciferin substrate for 4 hours before analysis. Luminescence was measured in 10 min intervals with Centro LB963 Luminometer (Berthold Technologies, Germany). Seventeen to 18 leaf discs per construct combination were analysed. The experiment was repeated twice.

### Protein production, purification, characterization and antibody preparation

For recombinant production of His_6_-tagged StTGA2.1, *Escherichia coli* BL21(DE3) cells were transformed with the pMCSG7 vector^77^ carrying the *StTGA2.1* cds, grown overnight and subsequently transferred to the liquid auto-induction media^100^, where they were incubated for 4 h at 37 °C and further 20 h at 20 °C to produce the protein. Cells were harvested by centrifugation, lysed and the protein purified by nickel affinity chromatography using the His- Trap HP column coupled with size-exclusion chromatography (SEC) using the HiPrep 26/60 Sephacryl S-200 column (GE Healthcare Life Sciences, UK). The protein was eluted into a buffer containing 30 mM Tris, pH 7.5, and 400 mM NaCl, and used for rabbit polyclonal anti- StTGA2.1 antibody preparation, provided by GenScript (USA).

The protein oligomeric state was determined based on SEC elution volume and Gel Filtration LWM Calibration Kit (standard sizes: conalbumin 75 kDa, ovalbumin 44 kDa, carbonic anhydrase 29 kDa, ribonuclease A 13.7 kDa and aprotinin 6.5 kDa, GE Healthcare, USA). Additionally, the His_6_-tag was removed by His_6_-tagged TEV protease cleavage and a secondary nickel affinity chromatography followed by an additional SEC, as well as an anion-exchange chromatography purification step. Chemical crosslinking was performed after His_6_- tag removal, for which the protein buffer was exchanged to 30 mM Hepes, pH 7.5, 400 mM NaCl using ultrafiltration with Amicon Ultra centrifugal filter units (Merck, Germany). The reaction was performed using the BS^3^ crosslinker according to the manufacturer’s instructions (Thermo Scientific, USA) and the protein oligomeric state evaluated by SDS-PAGE.

The *E. coli* cell-free protein synthesis (CFPS) was used for the production of StTGA2.3. All CFPS reactions (total volume 30 or 75 μL) were performed as described previously^79^, with 20-24h incubation at 16, 20 or 25 °C. Either the empty pEPQD0KN0025 vector (Addgene plasmid #162283)^79^ or water was added to the reagent mixture to prepare a CFPS components reference. Proteins were detected by SDS-PAGE and Western blot, using anti-StTGA2.1 antibodies (diluted 1:4.000, GenScript, USA). Additionally, the protein identity was confirmed with mass spectrometry, performed at the Department of Biochemistry and Molecular and Structural Biology at the Jožef Stefan Institute (Slovenia).

### Surface plasmon resonance

Surface plasmon resonance measurements were performed on Biacore T200 (GE Healthcare, USA) at 25 °C at the Infrastructural Centre for Analysis of Molecular Interactions, University of Ljubljana (Slovenia). To prepare the DNA, the PRX07p_1, PRX07p_2, PRX15p_1 and PRX46p complementary primers (Integrated DNA Technologies, Belgium, Supplementary Table 13) were mixed in a 2:3 molar ratio (long:short primers) and annealed by cooling the mixtures from 95 °C to 4 °C. The resulting DNA fragments carried the selected 20 bp promoter regions with a 15-nucleotide overhang that allowed hybridization with the complementary biotinylated S1 primer^101^, immobilized on the streptavidin sensor chip (GE Healthcare, USA). StTGA2.3 protein-DNA binding experiments were performed in a running buffer containing 25 mM Tris, pH 7.4, 140 mM NaCl, 1 mM MgCl_2_ and 0.005% P20. For StTGA2.1 the running buffer contained 180 mM NaCl. Flow cell 1 was used as a reference and the DNA fragments were injected across the flow cell 2 at a flow rate of 5 µL/min to immobilize ∼50 response units.

A kinetic titration approach was used to study the interactions between the CFPS-produced StTGA2.3 protein, the CFPS components reference that lacked StTGA2.3 or the His_6_-tagged StTGA2.1 (18.75, 37.5, 75, 150, or 300 µM) and the DNA fragments. The highest concentration of total protein (264 µg/mL) and four sequential 1.5-fold dilutions were used for the CFPS-produced StTGA2.3 and the CFPS components reference. The proteins were injected across DNA at five concentrations, with no dissociation time between protein injections, at a flow rate of 30 µL/min. Regeneration of the sensor surface was performed with 50 mM NaOH solution for 10 s and 300 mM NaCl for 10 s at a flow rate of 30 µL/min. The sensorgrams for the StTGA2.3 or the StTGA2.1 proteins were double subtracted for the response of the reference flow cell 1 and for the response of the CFPS components reference or of the running buffer, respectively.

## ACKNOWLEDGEMENTS

We thank our colleagues Barbara Jaklič, Valentina Levak, Tjaša Mahkovec Povalej, Lidija Matičič, Nastja Marondini and Rebecca Vollmeier for technical support and laboratory assistance. We thank prof. Dr. Jim Haseloff and Dr. Fernán Federici (University of Cambridge, UK) for providing the plasmid containing H2B-RFP. Template to amplify N7 was a kind gift from Prof. Volker Lipka (Georg-August-Universität, Göttingen). This work was supported by the Slovenian Research Agency through the program P4-0165, the projects J4-1777 and J1-2467, the program 1000-21-0105 for young researchers, in accordance with agreement on (co) financing research activity in 2021, as well as by the European Union’s Horizon 2020 research and innovation program under grant agreement No 862858. Funding was also provided by the Plant-Microbe Interfaces (PMI) Scientific Focus Area in the Genomic Sciences Program of from the U.S. Department of Energy, Office of Science, Office of Biological and Environmental Research. This manuscript has been co-authored by UT-Battelle, LLC under contract no. DE-AC05-00OR22725 with the U.S. Department of Energy. The United States Government retains and the publisher, by accepting the article for publication, acknowledges that the United States Government retains a nonexclusive, paid-up, irrevocable, world-wide license to publish or reproduce the published form of this manuscript, or allow others to do so, for United States Government purposes. The Department of Energy will provide public access to these results of federally sponsored research in accordance with the DOE Public Access Plan (http://energy.gov/downloads/doe-public-access-plan, last accessed September 16, 2020).

