## Supplementary Figures and Tables for "A mini-TGA protein, lacking a functional DNA-binding domain, modulates gene expression through heterogeneous association with transcription factors"

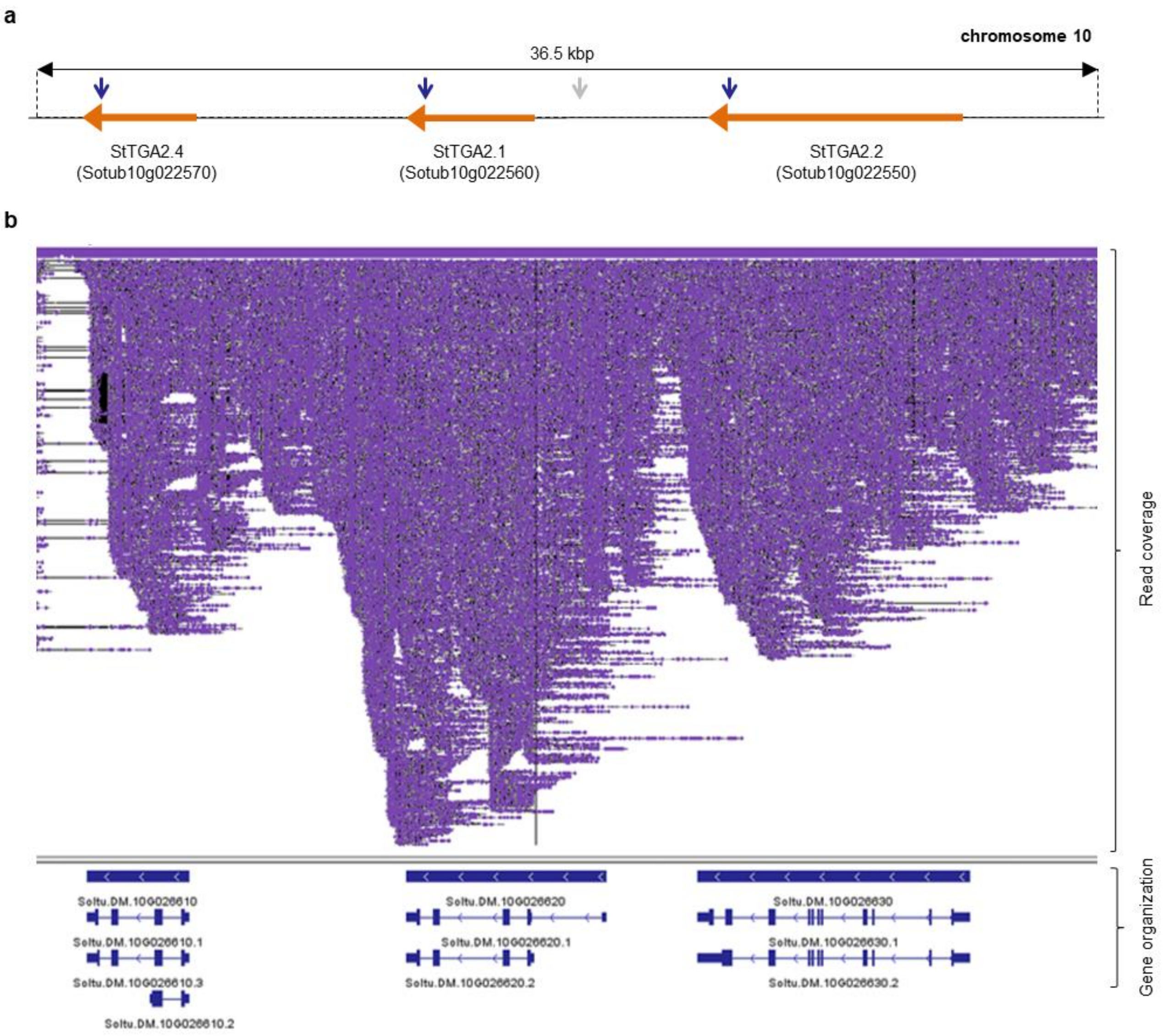


**Supplementary Fig. 1. Targeted long-read sequencing confirms the presence of mini-TGAs in potato genome. a,** Schematic representation of a ~36.5 kbp region on chromosome 10 in the double-monoploid (DM) potato reference genome v6.1[1], encompassing the three tandemly repeated StTGA genes, *StTGA2.1*, *StTGA2.2* and *StTGA2.4* (ROI). All three genes are located on the negative strand. Approximate annealing sites of primer pairs used in targeted long-read sequencing are indicated, with primer pair A (blue arrows) targeting all three genes, and primer pair B (grey arrow) targeting a central part of the ROI. The primers are listed in Supplementary Table 11. **b,** Screenshot, showing the read mapping (purple), obtained through targeted long-read sequencing of the ROI in the tetraploid potato cultivar Rywal, to the potato reference genome v6.1[1]. Complete sequence coverage (mean depth 569 at 99.6 % coverage of the ROI) confirms the presence and organization of the three StTGAs. Gene organization with several predicted alternative splicing variants, as represented in the DM genome with corresponding transcript IDs, is depicted at the bottom of the figure.

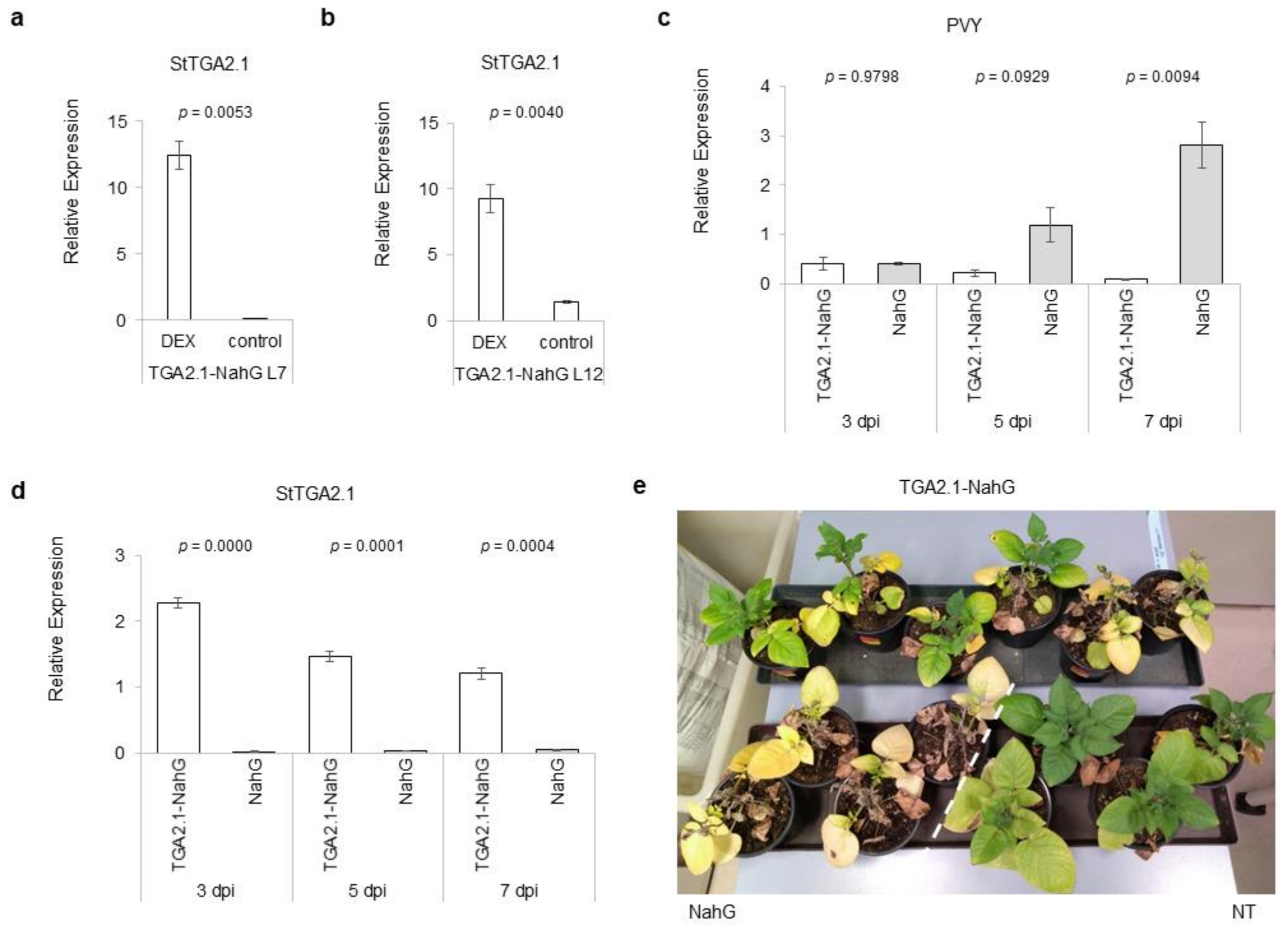


**Supplementary Fig. 2. PVY replication in the second salicylic acid-deficient transgenic line overexpressing *StTGA2.1*.** Relative expression levels of *StTGA2.1* in two NahG transgenic lines overexpressing *StTGA2.1* (TGA2.1-NahG), **a,** line 7 (L7) and **b,** line 12 (L12), three hours after dexamethasone (DEX) treatment, compared to non-treated plants (control). Average values ± standard error from three biological replicates for DEX treatment and **a,** two or **b,** three replicates for control are shown. Relative expression levels of **c,** PVY and **d,** *StTGA2.1* in PVY-infected leaves of DEX-treated TGA2.1-NahG (white) and NahG (light grey) plants at 3, 5 and 7 days post infection (dpi). Average values ± standard error from three biological replicates are shown. **a – d,** Significance was determined using a two-tailed *t*-test. **e,** Phenotypic differences in TGA2.1-NahG, NahG and non-transgenic potato (NT) plants at 32 dpi. NahG and NT plants are delineated with a white dotted line.

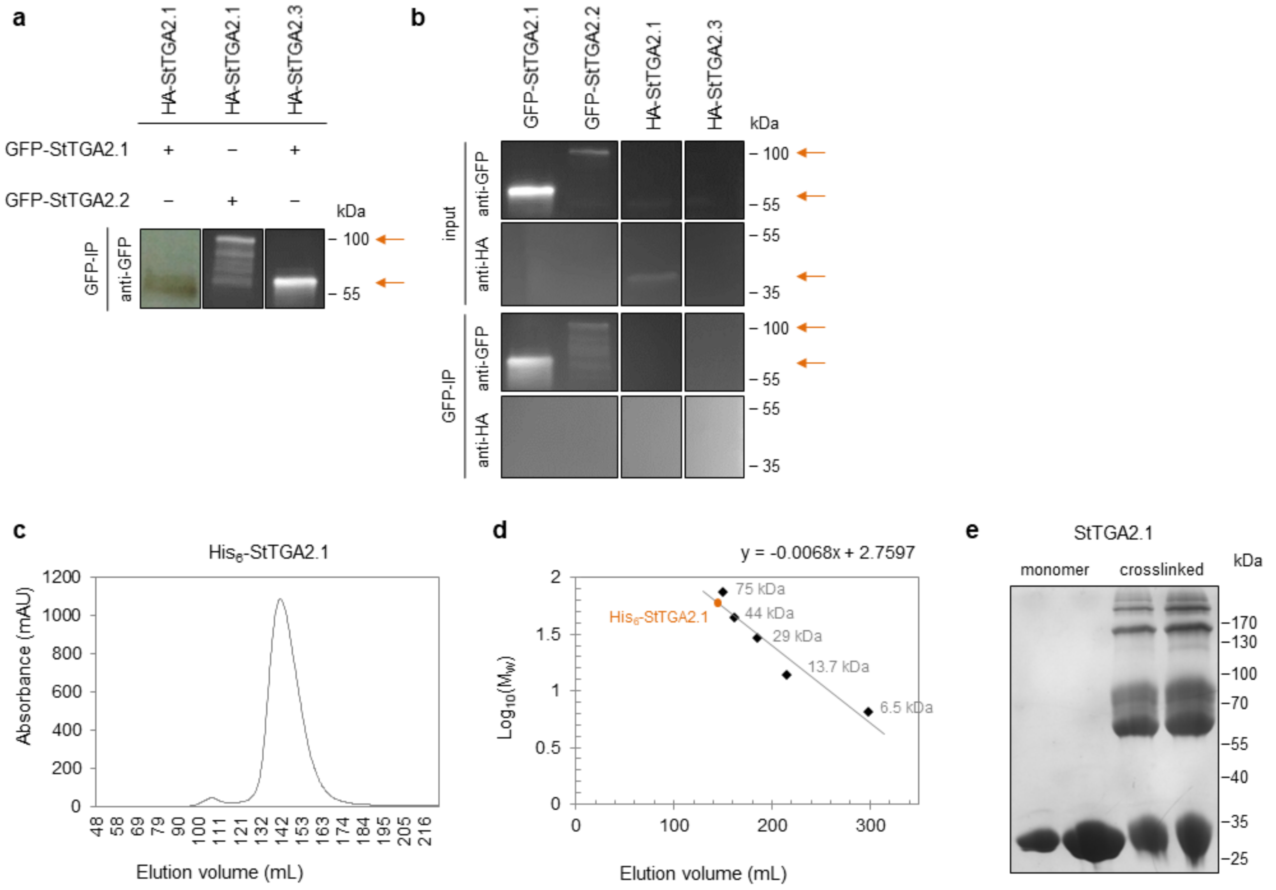


**Supplementary Fig. 3. Protein interaction analysis shows the mini-TGA StTGA2.1 can form homodimers *in planta* and *in vitro*. a,** Co-immunoprecipitation assay controls, showing the detection of GFP-tagged StTGAs after immunoprecipitation (GFP-IP) in the protein-interaction analysis samples (Fig. 3b). The combination of GFP and HA-tagged proteins expressed in *N. benthamiana* is indicated for each sample (+/−). **b,** Additional controls, showing individual GFP-tagged StTGAs could be detected in the leaf protein extracts (input) and GFP-IP samples. The HA-tagged StTGA2.1 alone was detected in the input, while none of the individual HA-tagged StTGAs were detected in after GFP-IP. **a,b,** Arrows indicate expected bands. **c,** Size exclusion chromatography elution volume of a purified His$_6$-tagged StTGA2.1. **d,** The protein standard calibration curve used to calculate its oligomeric state. The StTGA2.1 elution volume (orange dot) in relation to the protein standard (black diamonds) corresponds to an ~63 kDa large protein, which is twice the size of a monomer (~33 kDa). Protein sizes are depicted next to each standard marker. **e,** StTGA2.1 oligomeric state on SDS-PAGE gel, before (monomer) and after crosslinking (crosslinked), showing the formation of homodimers at ~60 kDa and higher order complexes at ~90, ~160 kDa and above. Two protein amounts per treatment were loaded.

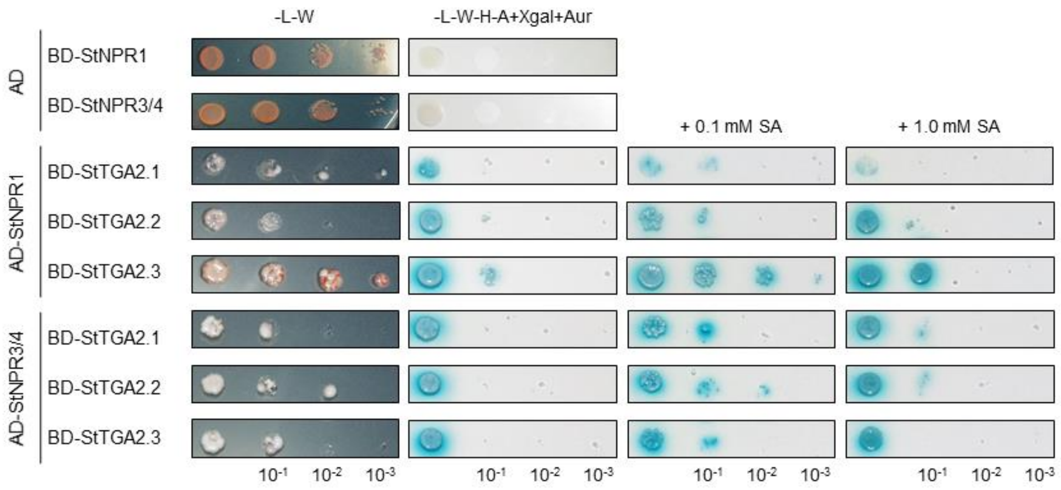


**Supplementary Fig. 4. Protein interactions between StTGAs and StNPR cofactors in yeast.** StTGA2.1, StTGA2.2, and StTGA2.3 interactions with StNPR1 and StNPR3/4 in the yeast two-hybrid assay. Yeast were co-transformed with bait (BD) and prey (AD) construct combinations and selected on control media without Leu and Trp (-L-W). Positive interactions were determined by yeast growth on selection media without Leu, Trp, His and Ade, with added X-α-galactosidase and Aureobasidin A (-L-W-H-A+Xgal+Aur). The effect of salicylic acid (SA) on interaction strength was determined by yeast growth on the same selection media with added 0.1 mM or 1.0 mM SA.

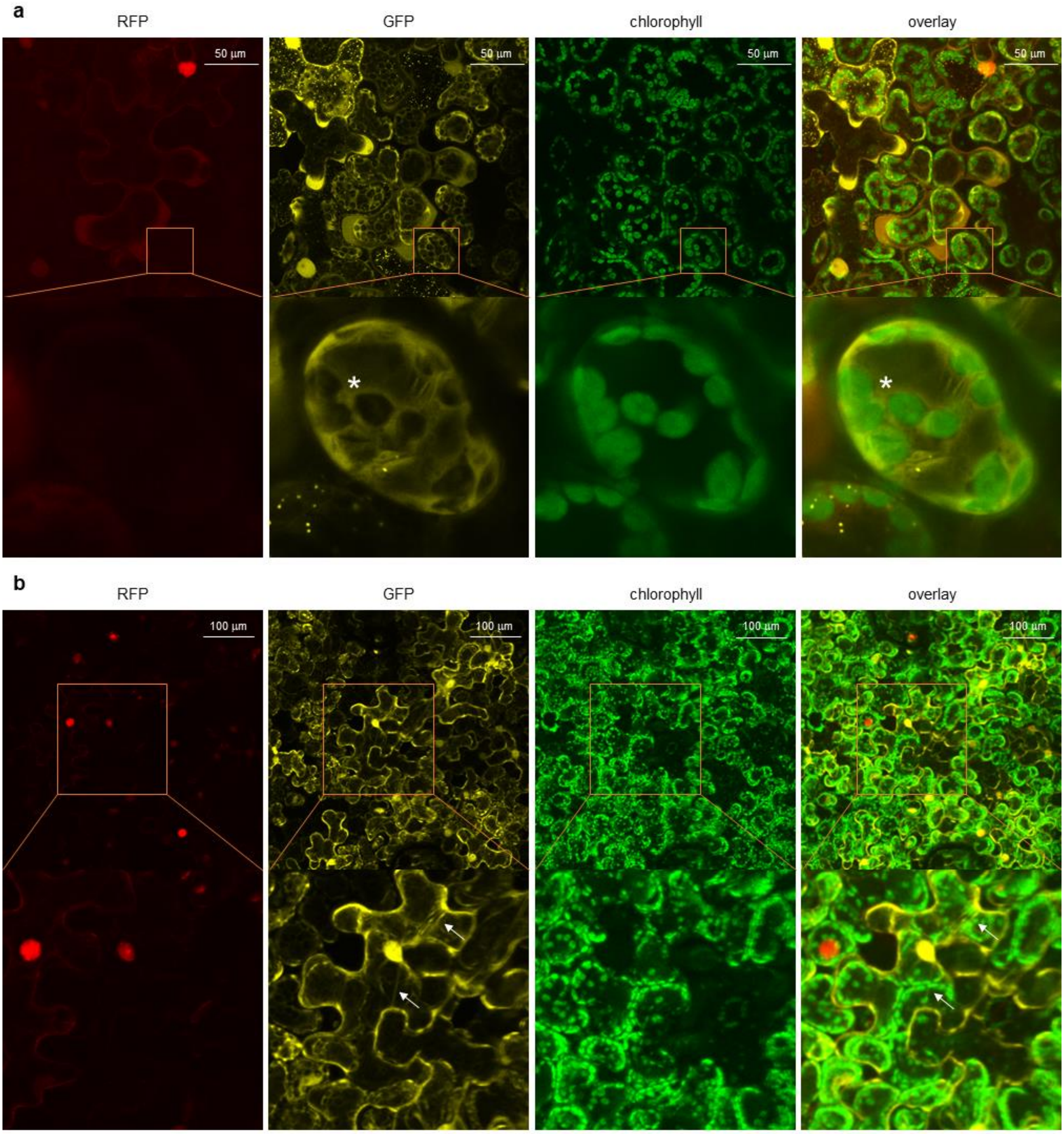


**Supplementary Fig. 5. Diverse localization patterns of StTGA2.1.** Subcellular localization of GFP-tagged StTGA2.1 (yellow) with H2B-RFP nuclear marker (red) and chloroplast autofluorescence (green) in *N. benthamiana* leaves, showing **a,** StTGA2.1 enrichment around chloroplasts (asterisk) and **b,** localization in the ER (white arrows). **a, b,** Protein fluorescence is represented as the z-stack maximum projection. Orange lines depict the image section close-up. Scale bars, **a,** 50 µm and **b,** 100 µm.

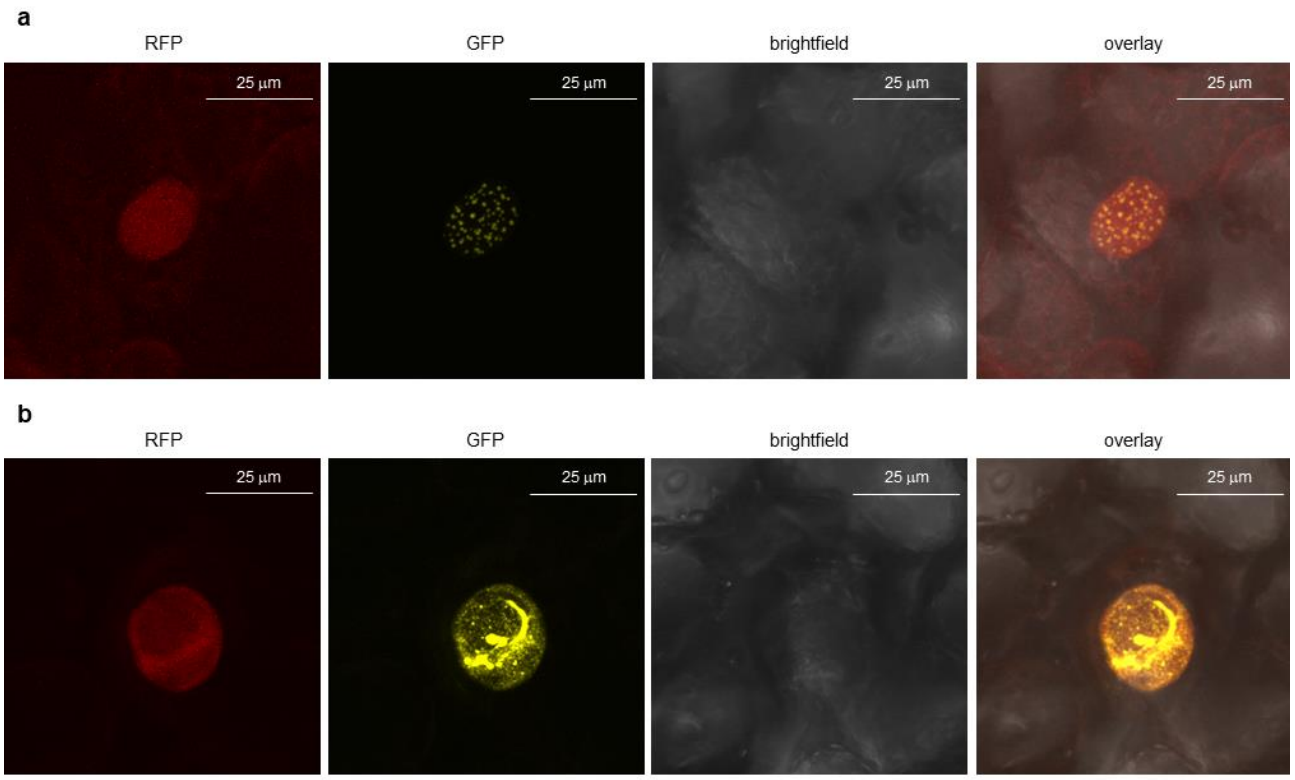


**Supplementary Fig. 6. StTGA2.2 and StTGA2.3 subnuclear formations.** Subnuclear localization of GFP-tagged **a,** StTGA2.2 and **b,** StTGA2.3 (yellow) with H2B-RFP nuclear marker (red) in *N. benthamiana* leaves. **a, b,** Protein fluorescence is represented as the z-stack maximum projection. Scale bars, 25 µm.

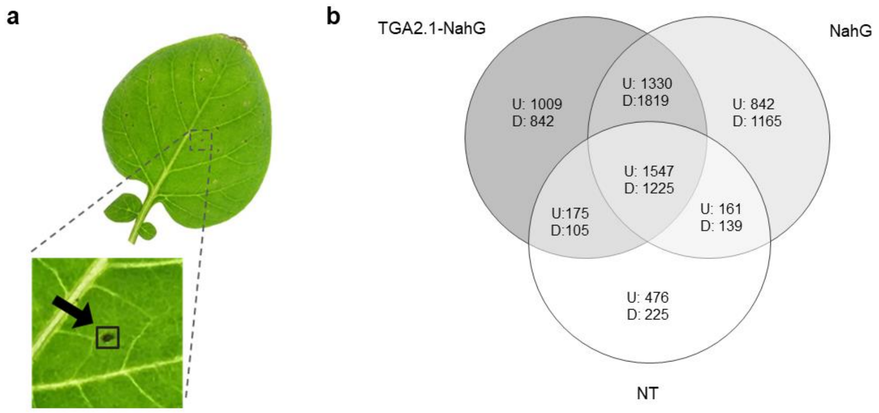


**Supplementary Fig. 7. RNA-sequencing sampling procedure and gene expression analysis Venn diagram. a,** Lesions with their immediate surrounding area were cut-out from PVY-inoculated leaves at four days post infection and pooled for RNA-sequencing analysis. Mock-inoculated leaf sections of similar size were sampled as controls. **b,** Venn diagram of differentially expressed genes from RNA sequencing between PVY and mock-inoculated TGA2.1-NahG, NahG and NT plants. Genes with adjusted p-value $< 0.05$ and $|\log_2 FC| \leq -1$ were considered significantly differentially expressed. U, up-regulated genes; D, down-regulated genes.

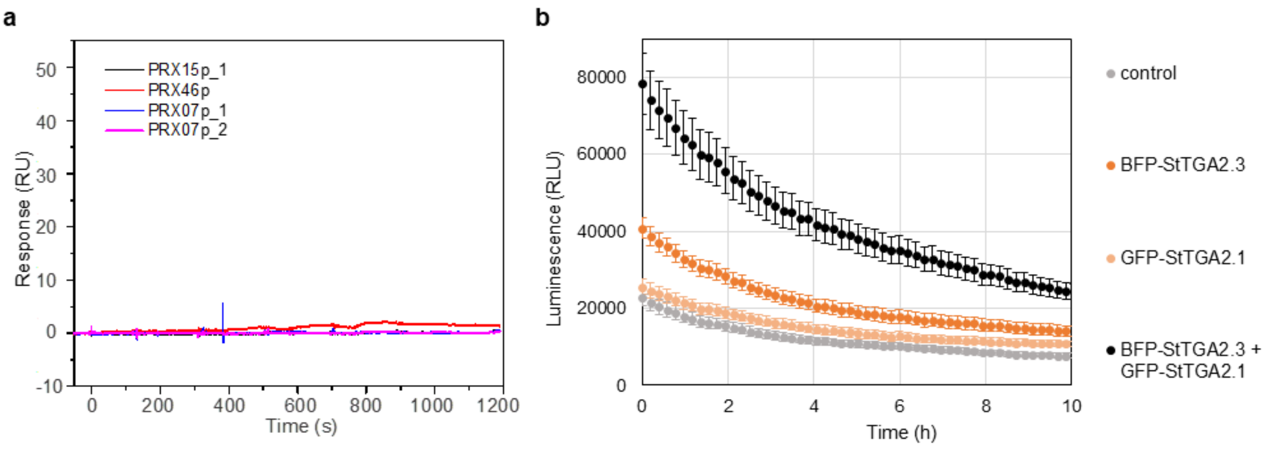


**Supplementary Fig. 8. Interaction between StTGA2.1 and selected TGA-binding sites and transactivation assay repetition. a,** Surface plasmon resonance results, showing no interaction between the StTGA2.1 protein and the chip-immobilized PRX15p_1, PRX46p, PRX07p_1 or PRX07p_2 DNA fragments, bound to the chip at ~77, 42, 55, or 60 response units (RU), respectively. Representative sensorgrams are shown. **b,** Transactivation assay repetition, showing *in planta StPRX07* promoter activation by GFP-tagged StTGA2.1 (light orange), BFP-tagged StTGA2.3 (dark orange) or a combination of both (black). BFP or GFP-tagged controls and their combination (control) were used to detect the basal promoter activity (grey). Average values ± standard error of 17 biological replicates in the first 10 h of measurement are shown.

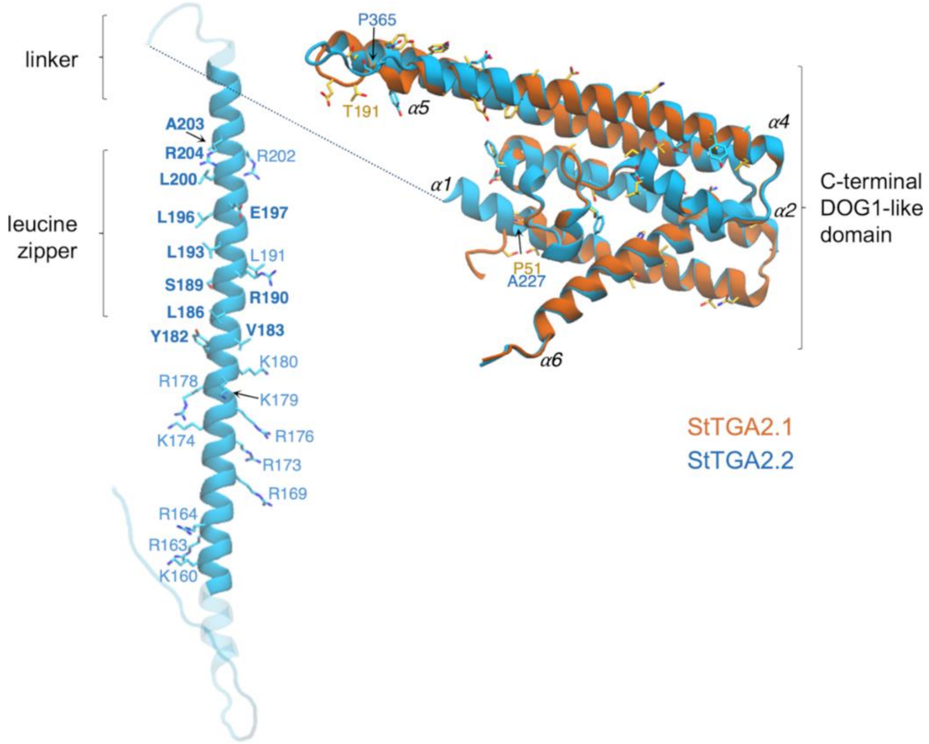


**Supplementary Fig. 9. Comparative structural analysis and persistent contacts in dimers of StTGA2.1 and StTGA2.2.** Molecular architectures of StTGA2.1 (orange) and StTGA2.2 (blue) proteins. The StTGA2.2 bZIP domain (aa 158-211) is shown. The N-termini of StTGA2.1 (aa 1-45) and StTGA2.2 (aa 1-129) are not shown for visual clarity. The C-terminal part is highly conserved between StTGA2.1 (aa 47-240), StTGA2.2 (aa 222-446), and StTGA2.3 (aa 105-327) (Fig. 5a). Amino acid residues forming persistent contacts in the leucine zipper, according to molecular dynamics simulations, are shown in bold. Basic amino acid residues that may contribute to DNA-binding are depicted and labelled. Non-conservative substitution sites in the putative DOG1 domains are also represented as liquorice.

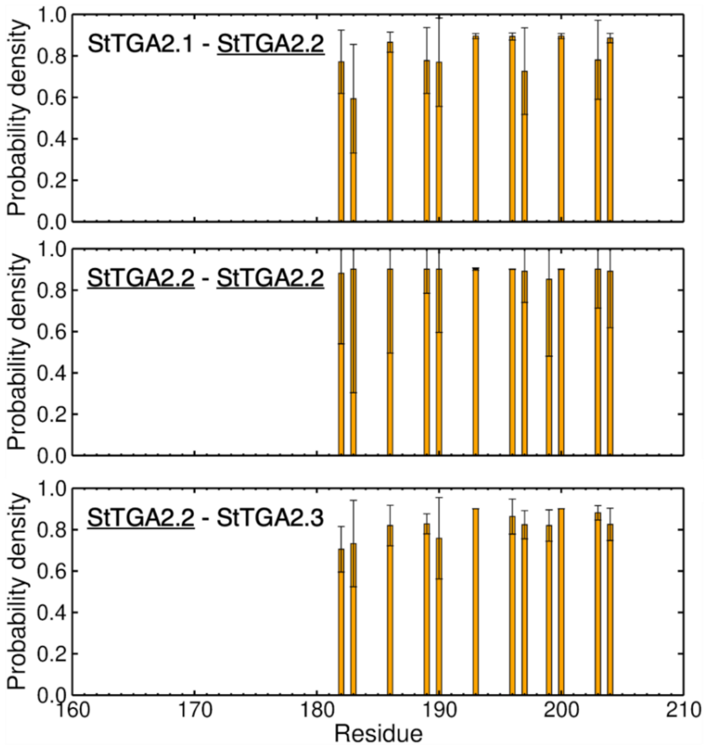


**Supplementary Fig. 10. Probability density of residues of StTGA2.2 forming contacts with a dimer partner.** The truncated protein forming a dimer with StTGA2.2 is specified in each plot (i.e., StTGA2.1, StTGA2.2, or StTGA2.3). A maximum distance of 7 Å between $C_\alpha$ atoms in a pair of residues was established. Bars with a standard deviation > 50% of the probability density are considered transient contacts in the simulations and are not included in these plots. In the StTGA2.2-StTGA2.2 and StTGA2.2-StTGA2.3 dimers the main interacting sites in StTGA2.2 are Tyr182, Val183, Leu186, Ser189, Arg190, Leu193, Leu196, Glu197, Leu200, Gln201, Ala203 and Arg204. The same interacting sites are kept in StTGA2.1-StTGA2.2, except for Glu197. In the simulations, StTGA2.1, StTGA2.2 and StTGA2.3, are truncated, keeping the amino acids 1-43, 159-206, and 42-89, respectively.

### Supplementary Tables

**Supplementary Table 1. A list of identified StTGA orthologues including basic protein information.** Protein sequence lengths, molecular weight ($M_W$) and theoretical pI, calculated with the ProtParam tool[2], and StTGA domain prediction based on Prosite[3].

| No. | Gene ID | Chromosome | Length (aa) | $M_W$ (kDa) | pI | Domain 1 | Domain 2 |
|---|---|---|---|---|---|---|---|
| 1 | **Sotub01g009430** | 1 | 327 | 36.34 | 8.61 | bZIP | DOG1 |
| 2 | **Sotub04g010500** | 4 | 369 | 41.66 | 6.20 | bZIP | DOG1 |
| 3 | **Sotub04g022350** | 4 | 369 | 41.57 | 5.46 | bZIP | DOG1 |
| 4 | **Sotub04g027470** | 4 | 361 | 40.88 | 6.68 | bZIP | DOG1 |
| 5 | **Sotub05g007640** | 5 | 483 | 53.10 | 6.51 | bZIP | DOG1 |
| 6 | **Sotub06g031310** | 6 | 503 | 55.94 | 6.53 | bZIP | DOG1 |
| 7 | **Sotub10g020240** | 10 | 488 | 55.53 | 6.26 | bZIP | DOG1 |
| 8 | **Sotub10g022140** | 10 | 461 | 52.24 | 6.47 | bZIP | DOG1 |
| 9 | **Sotub10g022550** | 10 | 438 | 48.23 | 7.18 | bZIP | DOG1 |
| 10 | **Sotub10g022560** | 10 | 270 | 30.49 | 5.69 | - | DOG1 |
| 11 | **Sotub10g022570** | 10 | 271 | 30.51 | 6.00 | - | DOG1 |
| 12 | **Sotub11g020650** | 11 | 324 | 36.17 | 8.86 | bZIP | DOG1 |
| 13 | **Sotub11g025820** | 11 | 435 | 48.60 | 6.91 | bZIP | DOG1 |
| 14 | **Sotub12g025350** | 12 | 348 | 39.56 | 6.92 | bZIP | DOG1 |

**Supplementary Table 2. Differential expression of StTGAs in NT and NahG genotypes after viral infection.** Microarray data showing StTGA gene expression comparisons between PVY- and mock-inoculated plants at 1, 3 and 6 days post inoculation (dpi), adapted from Baebler *et al*. (2014)[4]. Only statistically significant values (FDR adjusted p-value < 0.05) are shown, given as $log_2FC$. Cell shading based on $log_2FC$ values: blue, down-regulated; orange, up-regulated.

| | | NT | | | NahG | | |
|---|---|---|---|---|---|---|---|
| **Gene ID** | **Microarray ID** | **1 dpi** | **3 dpi** | **6 dpi** | **1 dpi** | **3 dpi** | **6 dpi** |
| **Sotub04g010500** | MICRO.14906.C1 | - | - | - | 1.04 | - | - |
| | MICRO.14906.C2 | - | - | - | 0.77 | - | -0.79 |
| | MICRO.14906.C3 | - | -0.93 | -0.68 | 0.49 | -0.90 | -1.35 |
| | POAED18TP | - | -0.45 | - | - | - | - |
| **Sotub04g022350** | MICRO.7564.C1 | 0.96 | - | - | - | - | - |
| **Sotub01g009430** | MICRO.14316.C1 | - | - | - | - | - | - |
| | MICRO.14867.C1 | 0.34 | 0.68 | 0.42 | 0.54 | - | 0.28 |
| | MICRO.8878.C1 | -0.62 | - | - | - | - | 0.53 |
| | POAC439TP | -0.64 | -0.39 | - | - | - | 0.60 |
| **Sotub10g022550** | MICRO.7441.C1 | -1.56 | -1.03 | 0.82 | -0.80 | 1.05 | 2.63 |
| | MICRO.7474.C3 | -0.95 | -1.06 | - | - | - | - |
| **Sotub10g022560** | bf_arrayxxx_0075b06.t7m.scf | - | - | - | - | 2.11 | 1.98 |
| | MICRO.7474.C2 | -0.41 | - | - | 0.34 | - | 0.80 |
| | MICRO.7474.C5 | - | - | - | - | 0.44 | 0.87 |
| | STMHL35TV | -0.90 | - | - | - | 0.70 | 1.72 |
| **Sotub10g022570** | MICRO.7474.C6 | 1.14 | 1.23 | - | 0.68 | - | - |
| **Sotub04g027470** | bf_arrayxxx_0086h05.t7m.scf | 0.82 | - | - | - | - | - |
| | MICRO.11212.C1 | - | - | -0.41 | - | -0.46 | -0.66 |
| | MICRO.9918.C1 | -0.37 | -0.46 | - | - | -0.48 | -0.47 |
| | STMGM64TH | -0.91 | -1.53 | - | -1.00 | - | 0.44 |
| **Sotub06g031310** | bf_suspxxxx_0045g08.t7m.scf | - | - | - | - | - | 0.71 |
| **Sotub10g020240** | bf_ivrootxx_0054a07.t3m.scf | 0.49 | -0.59 | - | - | - | - |
| **Sotub11g025820** | MICRO.14778.C1 | 0.67 | 0.78 | 1.15 | 0.71 | 1.23 | 2.22 |
| | MICRO.3439.C1 | -0.99 | -1.29 | -0.57 | -0.99 | -0.41 | 0.35 |
| **Sotub05g007640** | MICRO.15058.C1 | - | - | - | - | - | -0.74 |
| | POAD811TP | - | - | - | - | - | - |
| | POAD811TV | - | - | - | - | - | - |

**Supplementary Table 3. Technical validation of RNA sequencing results with qPCR.** Comparison of five differentially expressed genes obtained from RNA sequencing results (RNA-Seq) with qPCR analysis for TGA2.1-NahG, NahG and NT plants after viral infection (PVY- vs. mock-inoculated plants comparison), given as $log_2FC$. Gene expression ratios with FDR adjusted p-value < 0.05 are underlined. Cell shading based on $log_2FC$ values: blue, down-regulated; orange, up-regulated.

| | | TGA2.1-NahG | | NahG | | NT | |
|---|---|---|---|---|---|---|---|
| **Gene Name** | **Gene ID** | **RNA-Seq** | **qPCR** | **RNA-Seq** | **qPCR** | **RNA-Seq** | **qPCR** |
| **StACX3** | Sotub10g008540 | 2.05 | 2.07 | 2.21 | 2.11 | 1.46 | 1.50 |
| **StCS** | Sotub01g027350 | 1.55 | 1.17 | 1.49 | 1.80 | 0.52 | 0.67 |
| **StPti5** | Sotub02g020180 | 4.37 | 3.85 | 5.36 | 8.25 | 2.14 | 5.33 |
| **StPRX28** | Sotub01g042120<br>Sotub02g011690 | 7.47 | 6.29 | 6.91 | 5.69 | 2.89 | 2.48 |
| **StTGA2.1** | Sotub10g022560 | -1.45 | -1.54 | 0.21 | 0.95 | -0.26 | -0.37 |

**Supplementary Table 4. Enrichment of differentially regulated genes in salicylic acid-deficient transgenic plants overexpressing StTGA2.1 after viral infection (available as an Excel file).** Gene Set Enrichment Analysis results table showing MapMan ontology functional gene groups (BINs)[5] enriched in up-regulated or down-regulated genes in TGA2.1-NahG, NahG or NT plants after PVY infection. Functional groups, significantly enriched (FDR corrected q-value < 0.05) in at least one of the three genotypes, are listed. (+), enriched in up-regulated genes; (−), enriched in down-regulated genes.

**Supplementary Table 5. Classification of selected potato peroxidases.** Genes from the MapMan peroxidase functional group (BIN 26.12)[5] were named and classified according to BLAST results in the RedoxiBase database[6]. All hits with E-value equal to zero or best hits with E-value above zero (*) are listed for each gene. The presence of secretory signal peptides was predicted with SignalP 5.0[7]. (+), signal peptide; (−), no signal peptide predicted; (+/−) signal peptide predicted in at least one of several sequences assigned to the same gene ID; bold, the three peroxidases included in target confirmation.

| Gene ID | BLAST Hits | Classification | Secretory signal Peptide |
|---|---|---|---|
| PGSC0003DMG400020437 | StPRX05, StPRX52, StPRX72 | class III peroxidase | − |
| PGSC0003DMG400025492 | StPRX72, StPRX52 | class III peroxidase | + |
| PGSC0003DMG400025621 | StPRX20, StPRX09, StPRX28, StPRX50 | class III peroxidase | − |
| PGSC0003DMG400026575 | StPRX19, StPRX02, StPRX15 | class III peroxidase | +/− |
| PGSC0003DMG400032147 | StPRX12, StPRX10 | class III peroxidase | + |
| Sotub01g006690 | StPRX20, StPRX09 | class III peroxidase | + |
| Sotub01g042120 | StPRX28, StPRX50 | class III peroxidase | + |
| Sotub02g022480 | StPRX72, StPRX52 | class III peroxidase | + |
| Sotub02g022490 | StPRX72, StPRX52 | class III peroxidase | + |
| Sotub02g023700 | StPRX05 | class III peroxidase | + |
| Sotub02g027910 | StPRX12, StPRX10 | class III peroxidase | − |
| Sotub02g031070 | StPRX19, StPRX02 | class III peroxidase | + |
| **Sotub02g035680** | **StPRX15** | **class III peroxidase** | **+** |
| Sotub02g037370 | StPRX19* | class III peroxidase | + |
| **Sotub03g007840** | **StPRX46** | **class III peroxidase** | **+** |
| Sotub03g010400 | StPRX19, StPRX02 | class III peroxidase | + |
| Sotub03g015450 | StPRX24 | class III peroxidase | + |
| Sotub04g026530 | StPRX14 | class III peroxidase | + |
| Sotub04g026540 | StPRX11, StPRX17 | class III peroxidase | + |
| Sotub04g026550 | StPRX11, StPRX17 | class III peroxidase | + |
| Sotub05g006810 | StPRX75 | class III peroxidase | − |
| Sotub05g024920 | StPRX47, StPRX16 | class III peroxidase | + |
| Sotub06g009790 | StPRX47, StPRX16 | class III peroxidase | + |
| Sotub06g032420 | StPRX71 | class III peroxidase | + |
| Sotub06g033210 | StPRX25 | class III peroxidase | + |
| Sotub08g006050 | StPRX27 | class III peroxidase | + |
| Sotub08g011590 | StAPx-R | ascorbate peroxidase related | − |
| Sotub08g017100 | StPRX40 | class III peroxidase | + |
| Sotub09g007100 | StPRX03, StPRX04 | class III peroxidase | + |
| **Sotub09g020950** | **StPRX07** | **class III peroxidase** | **+** |
| Sotub10g024760 | StPRX03 | class III peroxidase | + |
| Sotub11g009570 | StPRX58 | class III peroxidase | + |
| Sotub11g029720 | StPRX51* | class III peroxidase | + |

**Supplementary Table 6. Biological validation of RNA sequencing results with qPCR.** Relative expression of three class III peroxidase genes as obtained by RNA sequencing (RNA-Seq) and qPCR analyses for both TGA2.1-NahG lines, line 7 (L7) and line 12 (L12), NahG and NT plants after viral infection (PVY- vs. mock-inoculated plants comparison), given as $\log_2$FC. Gene expression ratios with FDR adjusted p-value < 0.05 are underlined. Cell shading based on $\log_2$FC values: blue, down-regulated; orange, up-regulated.

| | | TGA2.1-NahG | | | NahG | | NT | |
|---|---|---|---|---|---|---|---|---|
| **Gene Name** | **Gene ID** | **RNA-Seq** | **qPCR L7** | **qPCR L12** | **RNA-Seq** | **qPCR** | **RNA-Seq** | **qPCR** |
| **StPRX07** | Sotub09g020950 | <u>10.78</u> | <u>6.87</u> | <u>3.86</u> | <u>7.13</u> | 3.54 | 5.19 | <u>3.43</u> |
| **StPRX15** | Sotub02g035680 | <u>1.90</u> | 1.14 | <u>1.59</u> | -0.15 | 0.20 | <u>4.27</u> | <u>3.42</u> |
| **StPRX46** | Sotub03g007840 | <u>6.79</u> | <u>3.92</u> | <u>3.24</u> | <u>2.88</u> | <u>2.21</u> | <u>4.23</u> | <u>1.84</u> |
| **StTGA2.1** | Sotub10g022560 | <u>-1.45</u> | <u>-3.67</u> | 0.10 | 0.21 | -0.07 | -0.26 | -0.50 |

**Supplementary Table 7. Statistics of interactions between the StTGA2.2 homodimer and DNA in molecular dynamics simulations.** Total frequency of hydrogen bond, salt bridge interactions, and hydrophobic contacts (*) between the protomers of the StTGA2.2 homodimer and the two DNA strands (DNA1 and DNA2) in five independent molecular dynamics simulations of 200 ns. For hydrophobic contacts, a maximum distance of 4 Å between $C_\beta$ atoms in the protein and the methyl group in a thymine was established.

| StTGA2.2 | DNA1 | Frequency / % | | | | | StTGA2.2 | DNA2 | Frequency / % | | | | |
|---|---|---|---|---|---|---|---|---|---|---|---|---|---|
| K160 | A-8 | 19 | 6 | 8 | - | 25 | K160 | - | - | - | - | - | - |
| R163 | T-7 | 90 | - | 82 | 63 | 59 | R163 | T-7 | 33 | 34 | - | 20 | 41 |
| R163 | A3 | - | 28 | 20 | - | - | R163 | - | - | - | - | - | - |
| R164 | A-8 | 27 | 28 | 1 | 19 | - | R164 | T-7 | 36 | 18 | 19 | 10 | 1 |
| Q167 | T-7 | 11 | - | 22 | 20 | 17 | Q167 | T-6 | 11 | 5 | 14 | 8 | 10 |
| N168 | A3 | 4 | 15 | 9 | 7 | 7 | N168 | C3 | 30 | 29 | 19 | 22 | 16 |
| N168 | T-4 | 26 | 16 | 24 | 23 | 26 | - | - | - | - | - | - | - |
| R169 | T2 | 22 | 22 | 30 | 14 | 19 | R169 | T2 | 19 | 13 | 20 | 24 | 24 |
| A171* | T-4 | 18 | 20 | 38 | 21 | 24 | A171* | T-5 | 21 | 34 | 38 | 33 | 44 |
| A172* | T2 | 43 | 24 | 21 | 21 | 43 | A172* | T2 | 37 | 38 | 36 | 32 | 34 |
| R173 | C-1 | 22 | 65 | 17 | 32 | 24 | R173 | G1 | 39 | 9 | 9 | 12 | 20 |
| R173 | G1 | 12 | 35 | 16 | 17 | 12 | R173 | C-1 | - | 6 | 7 | 11 | 12 |
| S175 | T-4 | 78 | 80 | 54 | 59 | 63 | S175 | C-4 | 60 | 59 | 58 | 49 | 57 |
| R176 | G1 | 26 | 38 | 31 | 41 | 39 | R176 | G1 | 13 | 34 | 20 | 17 | 13 |
| R176 | - | - | - | - | - | - | R176 | C-1 | 22 | 24 | 22 | 24 | 27 |
| R178 | T-4 | 37 | 29 | 33 | 33 | 22 | R178 | C-4 | 59 | 45 | 26 | 41 | 35 |
| R178 | G-5 | 18 | - | 58 | 17 | 24 | R178 | T-5 | 13 | 14 | 28 | 24 | 22 |
| K179 | G-3 | 18 | 7 | 31 | 17 | 10 | K179 | T-3 | 30 | 29 | 33 | 35 | 38 |
| K180 | A-2 | 22 | 38 | 30 | 28 | 31 | K180 | A-2 | 37 | 48 | 30 | 33 | 37 |

**Supplementary Table 8. Statistics of interactions between the StTGA2.1 in heterodimer and DNA in molecular dynamics simulations.** Total frequency of hydrogen bond, salt bridge interactions, and hydrophobic contacts (*) between StTGA2.1 in the StTGA2.1-StTGA2.2 heterodimer and DNA in five independent molecular dynamics simulations of 200 ns. For hydrophobic contacts, a maximum distance of 4 Å between $C_\beta$ atoms in the protein and the methyl group in a thymine was established.

| StTGA2.1 | DNA | Frequency / % | | | | |
|---|---|---|---|---|---|---|
| R7 | G1 | - | 10 | - | 22 | - |
| R7 | T2 | 22 | 13 | 22 | 17 | 25 |
| R11 | G1 | 31 | 12 | 20 | 27 | 12 |
| R11 | C-1 | 13 | 15 | 39 | 13 | 12 |
| R20 | T-5 | 27 | 5 | 22 | 15 | 0 |
| R20 | C-4 | 28 | 24 | 33 | 37 | 32 |
| R24 | T-3 | 28 | 22 | 15 | 24 | 22 |
| A13* | T-3 | 12 | 32 | 6 | 6 | 27 |

**Supplementary Table 9. Molecular dynamics simulations-based protocol for structure refinement of free and DNA-bound TGA dimers.** In the equilibration phases, there is a gradual change in temperature, time step, and the position restraint potentials. Atom velocities are re-distributed using five different seed numbers to initiate Equilibration_6. With that, five independent trajectories are generated for conformational sampling. Applied potentials are either simple harmonic restraints (H) or flat-bottom potentials (FB). During and after Equilibration_5, the position restraint potentials are applied only to the residues forming the leucine heptads. "Backbone" refers to both the backbone atoms of TGA proteins and DNA.

| Simulation Phases | Position Restraints | | | | Time Step | Total Time | Temperature |
|---|---|---|---|---|---|---|---|
| | Backbone | Type | Side Chain | Type | | | |
| **Equilibration_1** | 1.000 | H | 0.100 | H | 1 fs | 2 ns | 100.15 K |
| **Equilibration_2** | 1.000 | H | 0.050 | H | 2 fs | 1 ns | 200.15 K |
| **Equilibration_3** | 1.000 | H | 0.025 | H | 2 fs | 1 ns | 250.15 K |
| **Equilibration_4** | 1.000 | H | 0.000 | H | 2 fs | 1 ns | 300.15 K |
| **Equilibration_5** | 0.500* | H | 0.000 | - | 2 fs | 4 ns | 320.15 K |
| **Equilibration_6** | 0.250* | H | 0.000 | - | 2 fs | 5 x 2 ns | 340.15 K |
| **Equilibration_7** | 0.125* | H | 0.000 | - | 4 fs | 5 x 12 ns | 360.15 K |
| **Equilibration_8** | 0.050* | H | 0.000 | - | 4 fs | 5 x 128 ns | 340.15 K |
| **Equilibration_9** | 0.025* | H | 0.000 | - | 4 fs | 5 x 64 ns | 320.15 K |
| **Equilibration_10** | 0.250* | FB | 0.000 | - | 4 fs | 5 x 100 ns | 298.15 K |
| **Sampling** | 0.000 | - | 0.000 | - | 4 fs | 5 x 128 ns | 298.15 K |

*Position restraints applied to Ca only.

**Supplementary Table 10. Primers used for cloning and sequencing (available as an Excel file).**

**Supplementary Table 11. Primers and probes used for qPCR analysis.** F, forward; R, reverse; P, probe.

| Gene Name | Gene Description | Gene ID | Primer/Probe Sequence (5'-3') | Amplicon Efficiency |
|---|---|---|---|---|
| **StACX3** | acyl-CoA oxidase | Sotub10g008540 | Baebler *et al*. (2014)[4] | |
| **StCS** | citrate synthase | Sotub01g027350 | | |
| **StPRX28*** | class III peroxidase | Sotub01g042120, Sotub02g011690 | | |
| **StPti5** | ethylene responsive factor | Sotub02g020180 | F: GCAAGAAATACAGAGGCGTACGA<br>R: CATACTCTCGCACCGTGTCTAG<br>P: FAM-TTCGGCAGCGTATTTT-NFQ | 93% |
| **StTGA2.1** | TGA transcription factor | Sotub10g022560 | F: ATCTCTTGCTACTGAAGGGTCAT<br>R: CCATAGCCATTGCCATCTGACTTAT<br>P: FAM-CCGGAGATGCAGCTTATAAT-NFQ | 90% |
| **StPRX07** | class III peroxidase | Sotub09g020950 (CLCdnDe14_116472, CLCdnRY2_25372, CLCdnRY3_29096, VdnDe5_126154)** | F: GTTGGAATCGCACATTGTAATTTT<br>R: AGCTGCTAATGTAGGATCCATAGTCTT<br>P: FAM-TCCAAGATAGGCTATCGCCCGTACCTG-Zen Iowa Black TM FQ | 92% |
| **StPRX15** | class III peroxidase | Sotub02g035680 (CLCdnDe10_14846, CLCdnRY1_7366, CLCdnRY10_8871, VdnDe1_213633, VdnDe1_213635, VdnDe6_46588)** | F: CACTACAAACAAACAGATCACTTCACCTA<br>R: GACGAAGCAGTCGTGGAAGAA<br>P: FAM-TACCGCCGCCGCCACCCT-Zen Iowa Black TM FQ | 80% |
| **StPRX46** | class III peroxidase | Sotub03g007840 (CLCdnDe11_47734, CLCdnRY1_4154, PBdnRY1_4594, VdnDe1_299904, VdnDe1_299906, VdnDe6_44388)** | F: CGGCGGTTGGGTCAGTT<br>R: GAATGAGAGATGCTCCCATTCG<br>P: FAM-TGGCGGAAGCTATTGCGAGGGA-Zen Iowa Black TM FQ | 91% |
| **StCOX1** | cytochrome oxidase | Sotub04g015050 | Weller *et al*. (2000)[12] | |
| **StEF-1** | elongation factor | Sotub06g010680 | Baebler *et al*. (2009)[13] | |
| **PVY Univ** | PVY coat protein | AJ390300 | Kogovšek *et al*. (2008)[14] | |

* Previously named POX[4].

** For qPCR assay design we used sequences obtained from cultivar Rywal and cultivar Désirée reference transcriptome[15] as template. Only transcripts, targeted by the assay, are listed.

#### Supplementary Table 12. Primers used for targeted genome sequencing.

| Primer Pair | Primer Name | Sequence (5'-3') |
|---|---|---|
| A | WO100007-NIB-1_7F | GAGGCAAAATACGTGAAGCTAAAAG |
| | WO100007-NIB-1_7R | AGAGCACGGGCTGATTGTCT |
| B | WO100007-1_3F | GCTACGCCTTTGCAGCATAT |
| | WO100007-1_3R | ACAGGAAAAGCAACTCTGGCT |

**Supplementary Table 13. Complementary primers used for preparation of promoter DNA fragments.**

| Gene Name and ID | Promoter Fragment | Primer Name | Sequence (5'-3') |
|---|---|---|---|
| **StPRX07** Sotub09g020950 | PRX07p_1 | per950p_1F | GTTACTACTCGAGCGTGTGCCCAACGTCACAATCC |
| | | per950p_1R | GGATTGTGACGTTGGGCACA |
| | PRX07p_2 | per950p_2F | GTTACTACTCGAGCGTTAGGGGTGACGTTTCCAAT |
| | | per950p_2R | ATTGGAAACGTCACCCCTAA |
| **StPRX15** Sotub02g035680 | PRX15p_1 | per680p_1F | GTTACTACTCGAGCGTTTAATAATGATGACATTTG |
| | | per680p_1R | CAAATGTCATCATTATTAAA |
| **StPRX46** Sotub03g007840 | PRX46p | per840p_1F | GTTACTACTCGAGCGTGAACCTGAGGTCAACCGTT |
| | | per840p_1R | AACGGTTGACCTCAGGTTCA |
